# The Molecular Landscape of Asian Breast Cancers Reveals Clinically Relevant Population-Specific Differences

**DOI:** 10.1101/2020.04.09.035055

**Authors:** Jia-Wern Pan, Muhammad Mamduh Ahmad Zabidi, Pei-Sze Ng, Mei-Yee Meng, Siti Norhidayu Hasan, Bethan Sandey, Stephen-John Sammut, Cheng-Har Yip, Pathmanathan Rajadurai, Oscar M. Rueda, Carlos Caldas, Suet-Feung Chin, Soo-Hwang Teo

## Abstract

Molecular profiles of breast cancer have contributed to an improved understanding of the disease, enabled development of molecular prognostic signatures to guide treatment decisions and unveiled new or more accurate therapeutic options for breast cancer patients. However, the extent to which differences in genetic, environmental and lifestyle factors influence molecular profiles in different populations remains poorly characterised, as relatively few large-scale molecular studies of breast tumours in non-Caucasian populations have hitherto been reported. Here, we present the molecular profiles of 560 Asian breast tumours and a comparative analysis of breast cancers arising in Asian and Caucasian women. Compared to the breast tumours in predominantly Caucasian women reported in TCGA and METABRIC, we show an increased prevalence of Her2-enriched molecular subtypes and higher prevalence of *TP53* somatic mutations in ER+ Asian breast tumours. Using gene expression and immunohistochemistry, we observed elevated immune scores in Asian breast tumours, suggesting potential clinical response to immune checkpoint inhibitors. Whilst Her2-subtype and enriched immune score are associated with improved survival, presence of *TP53* somatic mutations is associated with poorer survival in ER+ tumours. Taken together, these population differences unveil new opportunities to improve the understanding of this disease and lay the foundation for precision medicine in different populations.

## INTRODUCTION

Breast cancer is a heterogenous disease, where histopathological and clinico-demographic features guide treatment choices – for example, use of endocrine treatment in estrogen-receptor positive (ER+) tumours, use of HER2-targeted treatment in Her2-positive tumours and use of chemotherapy in women with poor prognostic features, such as high grade. The development of molecular classifiers based on gene expression^1,2^ has led to a better understanding of the molecular subtypes of breast cancer, and the development of molecular-based prognostic tools such as OncoTypeDx has led to improved clinical decision-making – for example, in determining the benefit of chemotherapy in node-negative, ER+ disease^3^. Recognising that breast cancer is a copy number driven disease^4^, we have improved the stratification of breast cancers by integrating copy number alterations (CNAs) and gene expression in the classification of 2,000 primary tumours from the Molecular Taxonomy of Breast Cancer International Consortium(METABRIC)^5^. The subtypes identified, designated as Integrative Clusters (IntClust), have very distinct molecular features, drivers, and clinical courses^6,7^.

The molecular characterisation of breast tumours has also enabled the development of new therapies, and the selection of patients for the appropriate therapies. Gene expression^8,9^, targeted sequencing^7^, whole exome sequencing (WES)^10–12^, and whole genome sequencing (WGS) have revealed the genomics drivers of breast cancer, uncovering hitherto unrecognised therapeutic possibilities. Furthermore, mutational analysis has identified tumour mutational burden^13^ and homologous recombination deficiency^14^ as potential biomarkers for response to immunotherapy and poly ADP ribose polymerase (PARP) inhibitors^15,16^, respectively.

Notably, the extent to which findings from these studies, conducted in predominantly women of European descent, can be applied to other populations where there are differences in genetic, lifestyle and environmental factors remains largely understudied. Triple negative breast cancer is more common in African American women and a recent genomic analysis of 194 tumours show an increased HR deficiency signature, pervasive *TP53* mutations, and greater structural variations, indicating a more aggressive biology^17^. Breast cancer in Asians tend to occur at a younger age, with a higher proportion of pre-menopausal, oestrogen receptor (ER) negative and human epidermal growth factor receptor 2 (HER2) receptor positive disease (reviewed in Yap et al. 2019^18^). A recent genomic analysis of 187 early-onset Asian breast cancers show a higher prevalence of *TP53* mutations and enrichment in immune signatures^19^, whereas analysis of 465 triple-negative Asian breast cancers demonstrated broad similarities in tumours of the same subtype arising in Asian and Caucasian women^9^.

Given the potential impact of tumour subtypes, candidate drivers, mutational signatures and immune profiles on treatment options for breast cancer patients, and the hitherto lack of detailed information in Asian breast cancer patients, we have performed WES, shallow WGS (sWGS), and RNA-sequencing (RNA-seq) of 560 breast tumours from a cohort of Asian patients in Malaysia and report the impact of population differences on the molecular profiles of breast cancer.

## RESULTS

### Study Population and Clinicopathological Characteristics

Primary tumour tissue and blood samples (including 3 matched bilateral and 1 matched primary-recurrence samples) were obtained from 560 consecutive female patients with breast cancer treated at the Subang Jaya Medical Centre (SJMC), Malaysia between 2012 and 2017 who were recruited to the Malaysian Breast Cancer (MyBrCa) study^20^. After excluding samples that failed quality checks, data was generated for RNA-Seq (n=527), WES (n=546), and sWGS (n=533). Compared to patients in The Cancer Genome Atlas(TCGA)^10^, our patients were younger, presented at later stages, and had higher proportions of ductal carcinoma and Her2-positivity (28.9% MyBrCa versus 19.3% TCGA when NAs are excluded; Supp. Table 1). The MyBrCa tumour cohort includes Malaysians from a number of different ethnicities, primarily Malaysian Chinese (89%), Malay (4.6%) and Malaysian Indian (3.4%) (Supp. Table 1), and is as a whole genetically similar to other East/Southeast Asian populations according to genotyping analysis (Ho et al., in review). A principal component analysis of our RNA-Seq data did not reveal any significant differences across different ethnicities (SFig.1).

### Asian Breast Tumours Exhibit Higher Prevalence of Her2-positive disease

Clustering of the tumour profiles of the MyBrCa samples according to IntClust^5^ and PAM50^2^ (Figure 1 and SFig. 2, respectively) showed a higher prevalence of the Her2-enriched molecular subtypes, particularly IntClust 5 (13.1% in MyBrCa versus 7.9% in Caucasians; Figure 1A) and the PAM50 Her2-enriched subtype (23.3% in MyBrCa versus 9.9% in Caucasians; SFig. 2A). We also observed a higher prevalence of ER-negative Integrative Cluster 4 (IntClust 4-; 9.1% in MyBrCa, 7.6% in other Asians versus 4.6% in Caucasians; Figure 1A).

**Figure 1:**
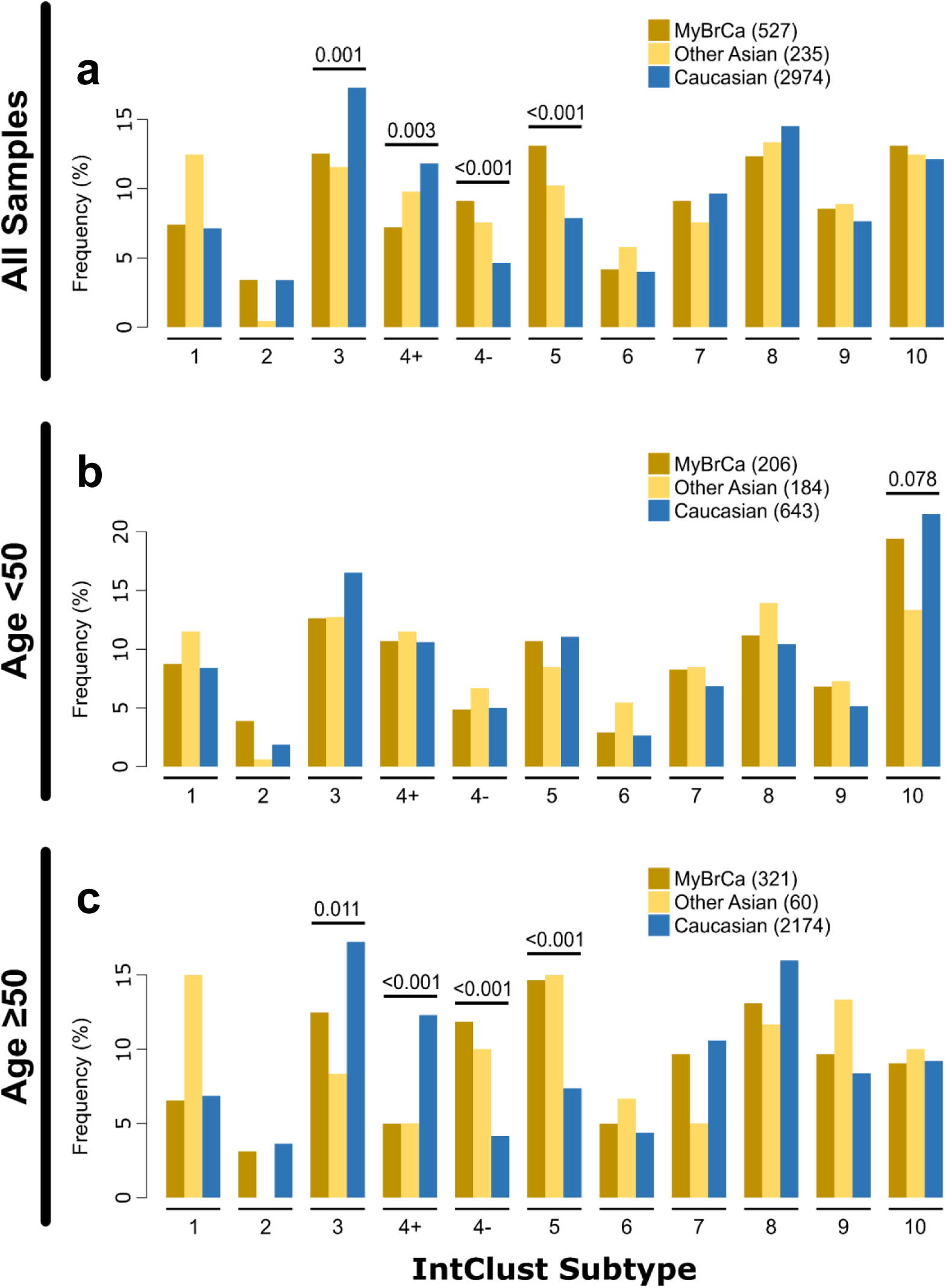
Molecular subtypes of Malaysian breast cancer. Comparison of Integrative Cluster molecular subtype distribution across Malaysian (MyBrCa), other Asian (Korean, TCGA Asian), and Caucasian (TCGA Caucasian, METABRIC, Nik-Zainal 2016) cohorts. Comparisons were done using the full cohorts **(a)** as well as with only patients below **(b)** or above **(c)** the age of 50. Numbers above the bars are p-values denoting significant differences between Asians and Caucasians for that subtype, as determined by Pearson’s chi-square test. Numbers in the figure legend indicate sample size.

Given that the median age of diagnosis of breast cancer was 49 years in Asians and 61 years in Caucasians, we examined the distribution of subtypes in women above and below the age of 50. In women below 50 years of age (n=206), there were no statistically significant differences between Asians and Caucasians in IntClust subtypes (Figure 1B), but there was an increased prevalence in the PAM50 Luminal B subtype (SFig. 2B). In women above 50 years of age (n=321; Figure 1C, SFig. 2C), we see the significant differences noted above in the whole population analysis, i.e. there was a markedly higher prevalence of Her2-positive disease: (a) IntClust 4- (11.8% in MyBrCa, 10% in other Asians, 4.1% in Caucasians), (b) IntClust 5 (14.6% in MyBrCa, 15% in other Asians, 7.4% in Caucasians), and (c) PAM50 Her2-enriched subtype (27.4% in MyBrCa, 30% in other Asians, 10.6% in Caucasians) suggesting that it is in this age group where these differences are mainly observed.

Finally, we also conducted a naïve combined cluster analysis using gene expression data from the MyBrCa and TCGA Caucasian cohorts in order to determine if there were any molecular subtypes that may be unique to Asians or Caucasians. However, unsupervised k-means consensus clustering did not reveal any exclusive clusters, but did support a higher prevalence of Her2-enriched subtypes and a lower prevalence of luminal subtypes in Asians (SFig. 3).

### Asian ER Positive Breast Tumours Exhibit Elevated TP53 Mutations

To understand the mutational architecture of MyBrCa tumours, we performed WES at a median coverage of 75X (range 5–123X) and 40X (range 5–77X) for 546 paired tumours and matched normal blood, respectively, and determined their somatic single nucleotide variant (SNV) and short insertion or deletion (indel) mutations. We identified 39,372 SNVs (18,508 non-synonymous) and 1,262 indels, with a median somatic mutation count of 45 SNVs and 2 indels per sample. We found similar subtype-specific non-synonymous sSNVs and indel mutations as previously reported (Fig. 2A)^7,21^. The most frequently mutated genes are *TP53* (42.9%), *PIK3CA* (27.8%), *GATA3* (9.7%), *MAP3K1* (5.5%), *KMT2C* (4.2%), *PTEN* (4.2%), *CBFB* (4.0%), *CDH1* (4.0%), *AKT1* (3.3%) and *NF1* (3.1%). Within the molecular subtypes, we observed that *TP53* mutation rates were high in IntClusts 5 and 10, and low in IntClusts 3 and 8. In contrast, *PIK3CA* mutation rates were high in IntClusts 3 and 8, with 96.4% of the *PIK3CA* variants being missense variants and concentrated at the known hotspot positions (SFig. 4). The majority of *GATA3* mutations were protein truncating (98.1%; SFig. 4), with high frequencies in IntClusts 1 and 8. *MAP3K1* mutation frequencies were high in IntClust 7, whilst mutations of *CBFB* and *CDH1* were high in IntClust 8.

**Figure 2:**
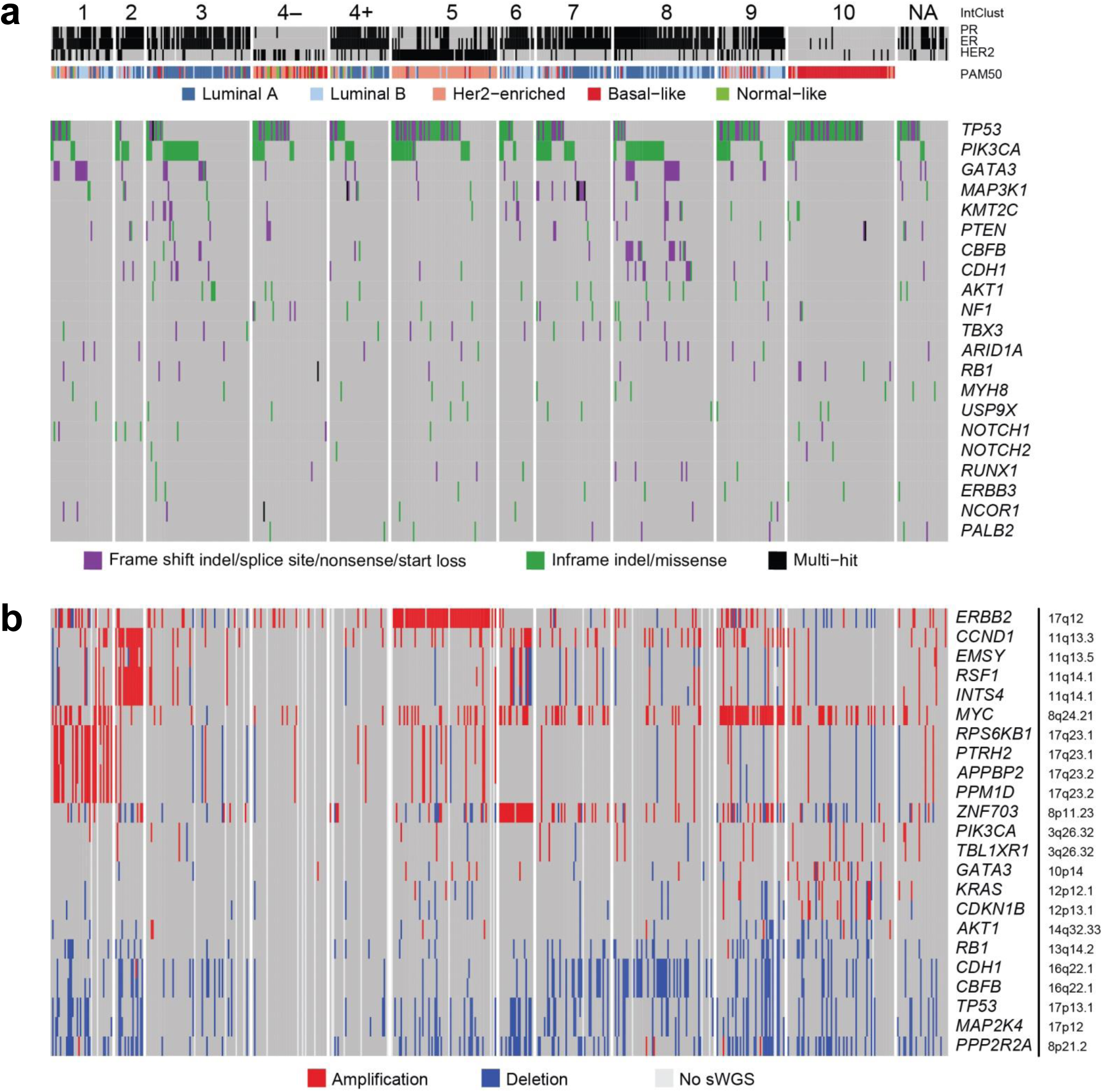
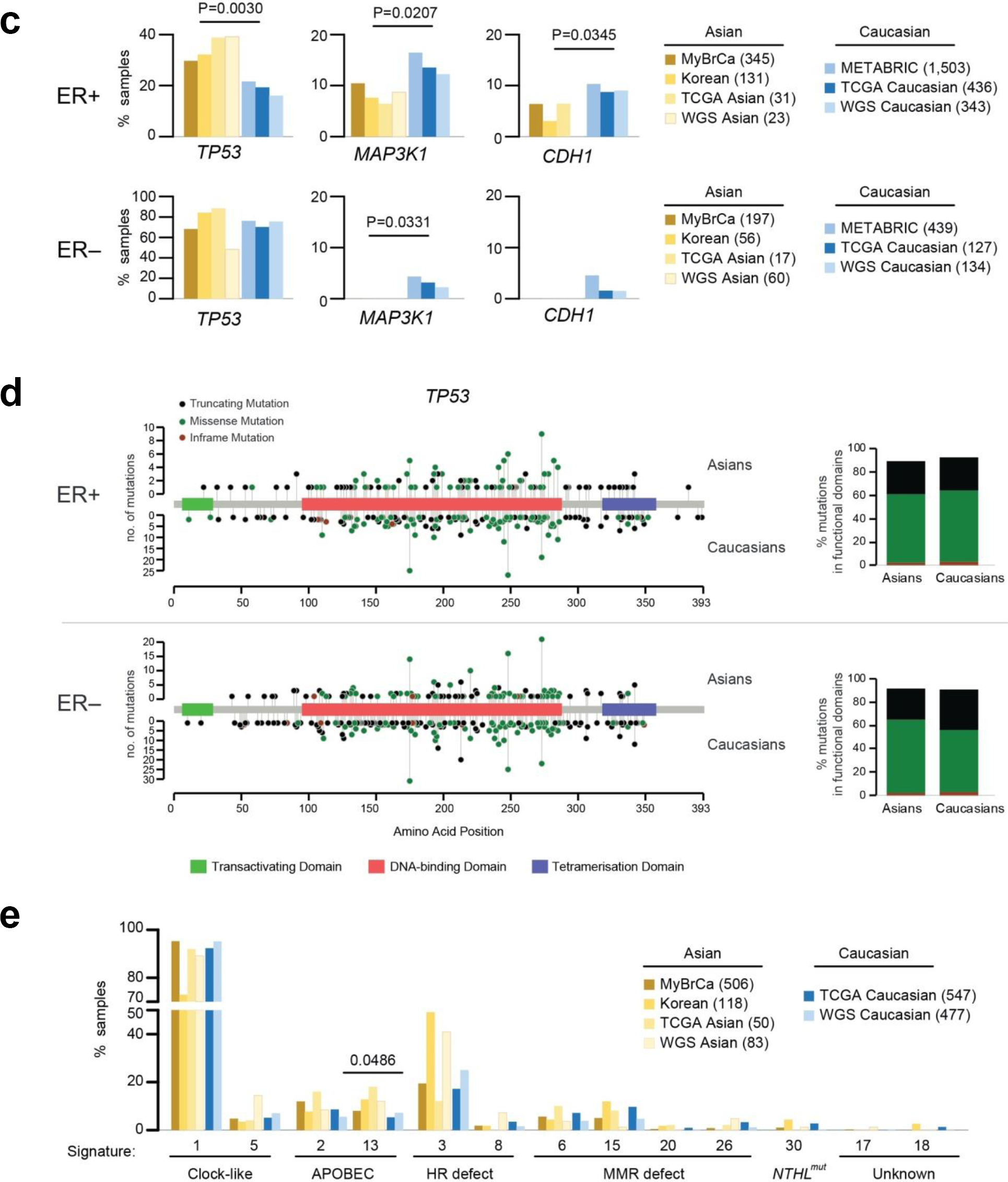
Mutational landscape of MyBrCa tumours. **a)** Somatic non-synonymous SNV and indel mutations of top mutated genes. Genes are sorted by the total mutation rates of MyBrCa cohort. **b)** Copy number aberration of breast cancer-related genes. Genes are sorted according to difference in number of samples carrying amplified over deleted copy of the genes, and genes within the same locus are grouped together for clarity. Samples are ordered as in **(a)**. NA: IntClust classification could not be determined due to unavailability of RNA-seq. **c)** Comparison of mutational prevalence of *TP53, MAP3K1*, and *CDH1* in Asian (MyBrCa, Kan et al., 2018 “Korean”, Asian TCGA, Korean samples Nik-Zainal et al., 2016 “WGS Asian”) and Caucasian (METABRIC, Caucasian TCGA, Nik-Zainal et al., 2016 “WGS Caucasian”) breast tumours, separated by ER status. **d)** Positions of *TP53* mutations in ER+ and ER– samples, separated by ethnicity. Right: Percent of mutations in each category that are located in transactivating, DNA-binding, or tetramisation domains of TP53. **e)** Frequencies of samples in Asian or Caucasian datasets carrying breast cancer-related mutational signatures. P-value from 2-sided student’s *t*-test.

Interestingly, *TP53* mutations were more common in Asians compared to Caucasians (42.9% compared to 30-35%; SFig. 5), and this difference was observed only in ER+ tumours and IntClust 8 (Fig. 2C, SFig 5-7; p=0.003 and p=0.035 respectively). However, there was no significant difference in the location of the mutations, with 83% and 84.8% of mutations occurring in the DNA binding domain of *TP53* for ER+ Asian and Caucasian samples, respectively (86.6% vs 83% for ER– samples; Fig 2D). *MAP3K1* mutation rates were lower in Asians irrespective of ER status (Fig 2C; p=0.02 and p=0.03 in ER+ and ER–, respectively), while *CDH1* mutation rates were lower in ER+ Asian tumours (p=0.03), although there was no difference in *CDH1* mutation rates after accounting for histological subtype (SFig. 8).

We performed driver gene analyses^22^ (SFig. 9, Supp. Tables 2 and 3) and identified well-established breast cancer oncogenes and tumour suppressors. For example, regardless of ER status or ethnicity, *PIK3CA* and *AKT1* have high oncogene (ONC) but low tumour suppressor gene (TSG) scores, consistent with their established roles as known breast cancer oncogenes, while known tumour suppressors *TP53, RB1* and *PTEN* have elevated TSG scores. We also observed high TSG scores for *MAP3K1* and high ONC for *CDKN1B* in ER+ as previously noted^7^. We also observed preferentially higher *SF3B1* mutations in ER+ samples, with the recurrent K700E mutation enriched in Caucasians ER+ tumours^23^ (Caucasian ER+ ONC=58.6 vs Asian ER+ ONC=12.5). We additionally examined pairwise association of somatic mutations (SFig. 10) in Asian or Caucasian breast tumours. We observed mutual exclusivity between *TP53* and each of *CDH1* and *GATA3*, between *PIK3CA* and each of *CDH1* and *MAP3K1*, and between *GATA3* and *CBFB* as previously noted^7^ in Asians and Caucasians tumours (SFig. 10). Finally, an analysis of tumours with germline pathogenic variants in 6 high to moderate breast cancer susceptibility genes did not reveal any significant differences between our cohort and TCGA^24^ (SFig. 11).

Analysis of copy number aberrations (CNA) using whole genome sequencing at 0.1X coverage revealed frequently altered regions in chromosomes 1, 8, 16, and 17 (SFig. 12), similar to what has been found in previous large genomic studies (TCGA, METABRIC). High level amplifications were observed in known oncogenes such as *ERBB2* and *MYC*, while deletions were observed in loci encompassing *TP53* and *MAP2K4.* We observed IntClust-specific CNAs (Fig. 2B and SFig. 13): for example, amplification of *ERBB2* at 17q12 in IntClust5; amplification of 17q23 encompassing *RPS6KB1, PTRH2, APPBP2* and *PPM1D* in IntClust1, amplification of two distinct amplicons around 11q13 (*CCND1* and *EMSY, RSF1*, and *INTS4*) in IntClust2, and amplification of *ZNF703* at 8p12 in IntClust6.

We also used copy number data to estimate the fraction of cancer cells that harbour mutations for nine common driver genes, and found that most genes had median mutational cancer cell fractions (CCF) of 1, suggesting that mutations in these genes are clonal and tend to occur early during tumour development (SFig. 14).

Together, WES and sWGS analyses suggest that the subtype specific somatic SNVs, indels, CNA and cancer cell fraction patterns in our cohort are similar to that previously described in other Asian or European populations, with the notable exception that *TP53* mutations are more common in ER+ Asian breast cancers.

### Asian and Caucasian Breast Tumours Exhibit Similar Mutational Signatures

We compared the mutational signatures previously detected in breast tumours^21,25,26^: Signature 1 and 5 (clock-like), Signatures 2 and 13 (APOBEC enzymatic activity), Signatures 3 and 8 (DNA repair deficiency), Signatures 6, 15, 20 and 26 (mismatch repair (MMR) deficiency), and Signature 30 (base-excision repair protein Nth Like DNA Glycosylase 1 (NTHL1) deficiency). We also included Signatures 17 and 18 that have been previously detected in breast tumours, yet of unknown aetiology.

We found only minor differences in the mutational signatures between Asian and Caucasian breast tumours (Fig. 2E and SFig. 15). Whilst we observed a higher percentage of Korean samples (Korean and WGS Asian) with the HR deficiency Signature 3 as previously reported^19^, this was not observed in other Asian datasets (MyBrCa and TCGA Asian; Fig. 2E). Significantly, we found a higher percentage of Asian breast tumours with Signature 13, consistent with the higher prevalence of the *APOBEC3B* germline deletion polymorphism among the population (p = 0.0486). No differences for other mutational signatures were observed (Fig. 2E).

We extended the analysis to molecular subtypes and found no significant difference in the distribution of mutational signatures based on ethnicity. IntClusts 3 and 8 samples primarily have high Signature 1, and moderate number of samples with Signatures 2, 13 and 3, and consistent with this, ER+ luminal A tumours by PAM50 classification show an elevated percentage of samples with Signature 1 as well (SFig. 16 and 17). IntClust 5 samples have high Signature 2 and 13, and consistent with this, the Her2-enriched tumours by PAM50 classification show an elevated percentage of samples with Signatures 2 and 13, suggesting increased APOBEC enzymatic activity^27^ within the Her2-enriched cohort. By contrast, IntClust 10 samples have a higher percentage of samples with the HR deficiency Signature 3, and consistent with this, the basal-like samples by PAM50 classification have a high percentage of samples with Signature 3. Together, these results suggest that there is no significant difference in the carcinogenic processes driving the development of each subtype of breast tumours in both Asian and Caucasian women.

### Asian Breast Cancers Exhibit Enriched Immune Scores

Given that population differences in the genetic, environmental and lifestyle exposures that drive carcinogenesis could shape the tumour microenvironment and lead to different outcomes, we performed pathway gene-set enrichment analyses (GSEA) to determine pathways that are differentially activated between our cohort and TCGA. Interestingly, we found that five out of the top fifteen most differentially expressed pathways were related to the immune system (Supp. Table 4). We determined the immune scores for MyBrCa, Korean, METABRIC and TCGA Caucasian cohorts according to five different scoring methods: ESTIMATE^28^, GSVA using gene sets for immune cells, GSVA using the expanded interferon-gamma gene set, the IMPRES method^29^, and CD8+ T-cells scores from CIBERSORT^30^. Remarkably, all five methods showed significantly elevated immune scores in both Asian cohorts relative to the Caucasian cohorts (Figure 3A).

**Figure 3:**
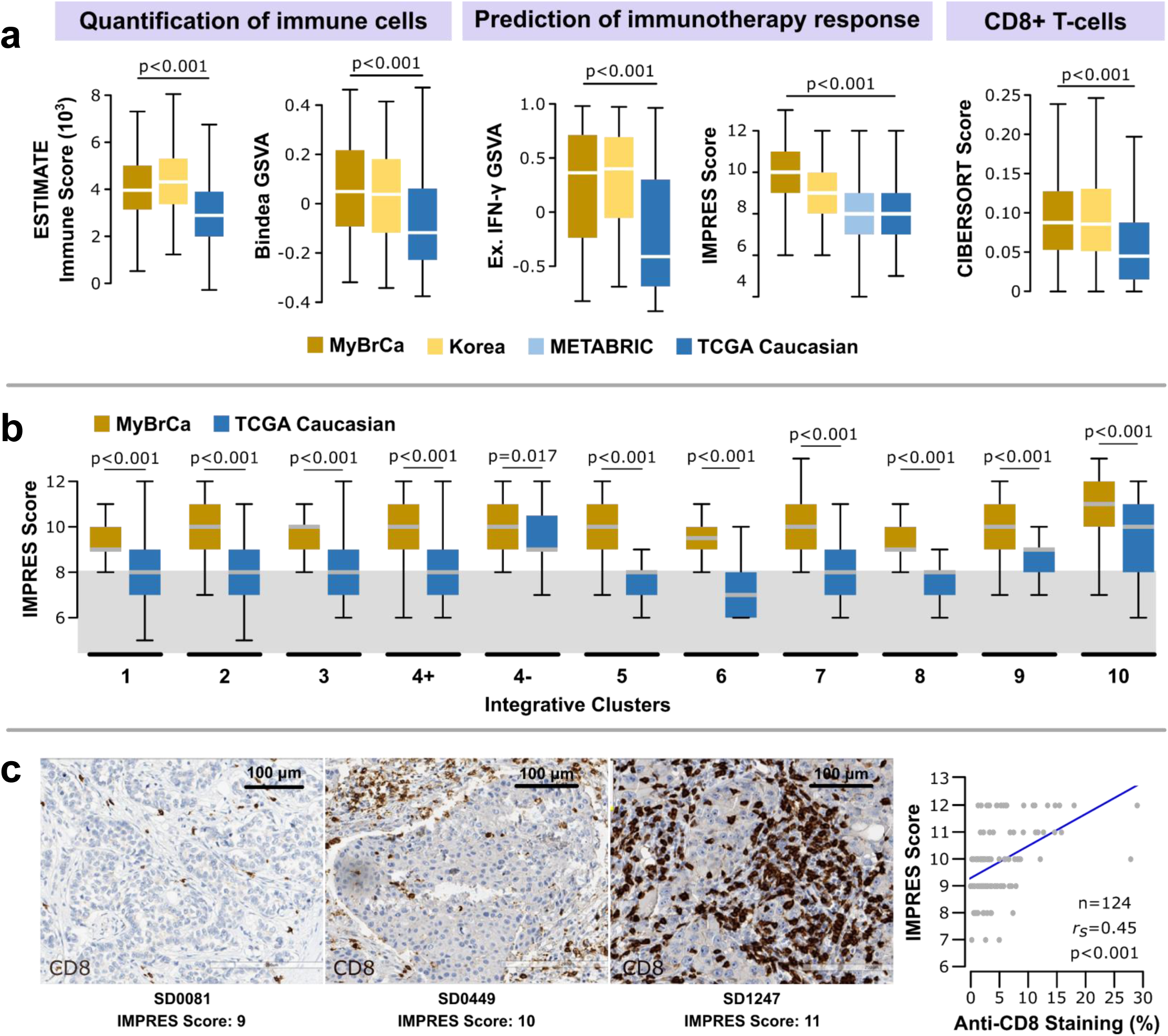
The tumour immune microenvironment of Malaysian breast cancer. **a)** Comparison of the tumour immune microenvironment in breast tumours from the Malaysian (MyBrCa, n=527), Korean (n=168), METABRIC (MB, n=997), and Caucasian TCGA (n=638) cohorts across five quantification methods – From left to right: ESTIMATE immune scores, GSVA using Bindea et al.’s (2013) combined immune gene set, GSVA using an expanded IFN-γ gene set (Ayers et al. 2017), IMPRES (Auslander et al. 2018), and CD8+ T-cell scores from CIBERSORT. Outliers not shown. **b)** Breakdown of IMPRES scores across SJMC and TCGA by IntClust subtype. Grey area indicates the threshold (score=8) used by Auslander et al. to separate high versus low scores. Outliers not shown. **c)** Validation of tumour immune scores in the Malaysian cohort using IHC; representative images shown with staining for CD8 in brown. Right: Correlation between percentage of cell stained with anti-CD8 by IHC versus IMPRES scores.

As immune profiles are subtype-dependent, we compared IMPRES scores between MyBrCa and the TCGA Caucasian cohort across breast cancer subtypes. We observed the highest immune enrichment was in IntClust 10, which comprises mainly of TNBCs, in concordance with published studies^31,32^. More interestingly, we found that the IMPRES scores were significantly higher for MyBrCa cohorts compared to TCGA Caucasian samples independent of subtype classification (Figure 3B). Similar patterns were found using the other immune scores (SFigure 18) and classification methods (SFigure 19), suggesting a systemic enrichment of immune scores in Asian breast cancers that is unrelated to their molecular subtype.

To identify immune cell types that drive the enrichment of immune scores in Asians, we used CIBERSORT on RNA-seq gene expression profiles to quantify the relative abundance of 22 different immune cell types in the tumour immune microenvironment. We found CD8+ T-cells and macrophages are enriched in Asian breast cancers (Figure 3A, SFigure 20), and this was consistent with TIMER and GSVA with Bindea gene sets (SFigure 21-22), suggesting an overall enrichment of cytotoxic NK and T cells in Asian tumours.

In a subset of 124 patients our cohort, we compared each patient’s IMPRES score with anti-CD8 staining of tumour-infiltrating lymphocytes (TILs) in archival formalin-fixed paraffin-embedded (FFPE) blocks and found a high correlation (r_s_ = 0.45; Figure 3C). Similarly, IMPRES scores were highly correlated with CD3 IHC scores (r_s_ = 0.51; SFigure 23), and ESTIMATE immune scores were highly correlated with CD3 and CD8 IHC scores (SFigure 23). Together, these data suggest that differences in immune scores derived from RNA-seq data are at least in part due to differences in the TILs within each tumour.

### Factors Contributing to Enriched Immune Scores

To explore the factors contributing to enriched immune scores in Asians, we first explored whether the generation of neo-antigens through different mutational processes is associated with immune scores. Indeed, we found that Signatures 2 and 13 (APOBEC) and Signature 3 (HR deficiency) are positively correlated with immune scores, whereas Signature 1 (aging) is negatively correlated (Fig. 4B).

**Figure 4:**
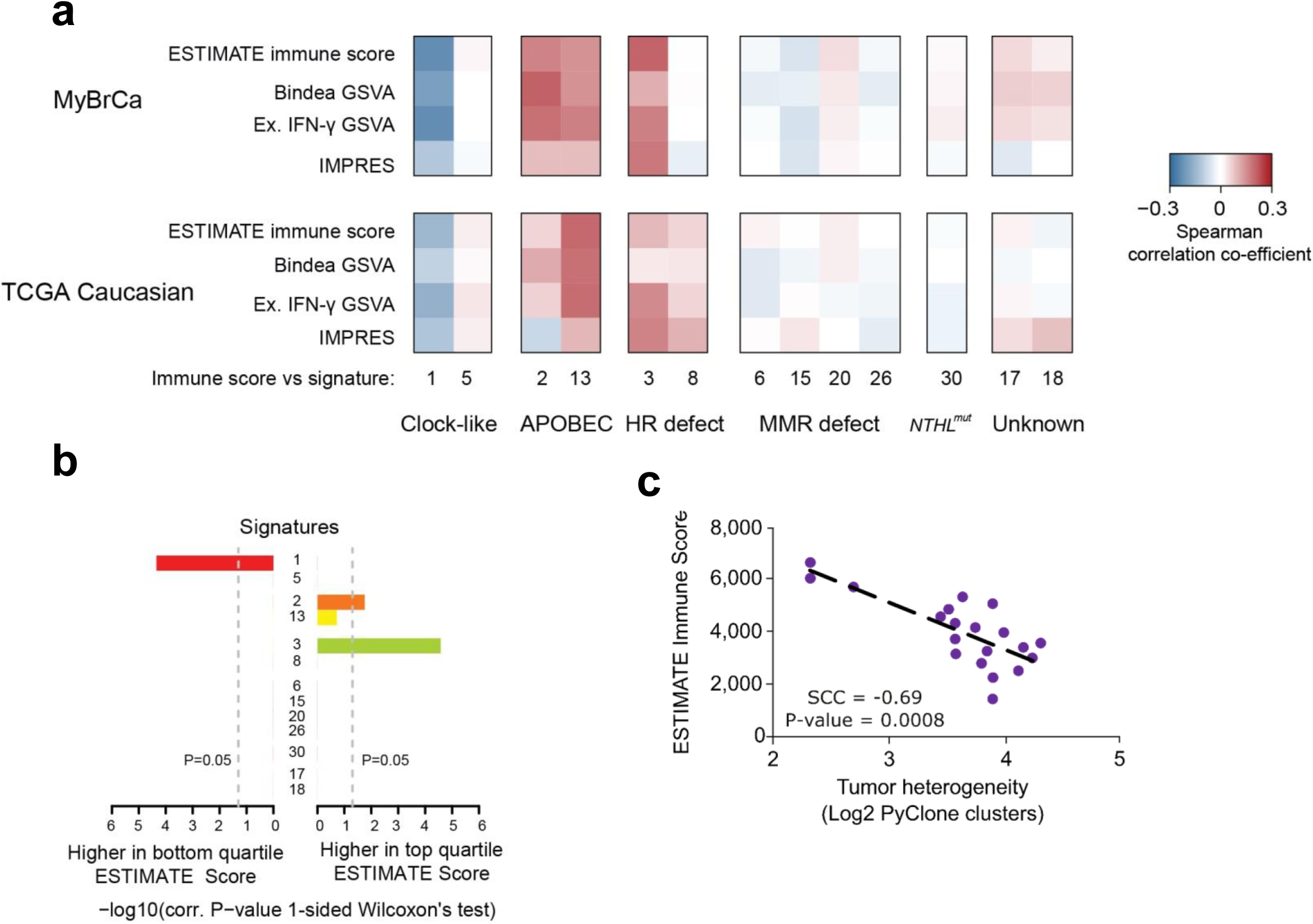
Factors associated with immune scores. **a)** Heatmap depicting Spearman correlation of co-efficient between mutation signatures in MyBrcA or TCGA Caucasian samples versus immune scores. **b)** Bar plots depicting P-values from 1-sided Wilcoxon’s test of the top versus bottom quartile MyBrCa samples, ranked according to ESTIMATE scores. **c)** Comparison of ESTIMATE immune scores to the tumour heterogeneity of each MyBrCa sample, quantified as the log_2_ of the number of clusters predicted by PyClone analysis. Samples were stratified into 20 groups according to ESTIMATE immune score.

We also asked if the immune signatures were associated with the underlying tumour heterogeneity of each sample. To quantify tumour heterogeneity, we used PyClone^33^ to estimate the number of subclonal clusters in each sample. A comparison of ESTIMATE immune scores with the number of PyClone clusters revealed a strong negative association (*r*_*s*_ = -0.69, p = 0.0008; Fig. 4D), suggesting that tumours in the MyBrCa cohort with high immune scores have correspondingly low tumour heterogeneity. Furthermore, we also found that the MyBrCa samples have significantly lower tumour heterogeneity relative to TCGA samples across subtypes (SFigure 24). These data are consistent with a model in which the stronger immune response in Asian breast cancer patients leads to higher selective pressure on tumour cells and, ultimately, more homogenous tumours in those patients^34,35^.

Next, in order to identify clinical or demographic factors that could contribute to the elevated immune scores observed in Asian breast cancer samples, we conducted linear regression analysis of immune scores against all available clinical and demographic data, including age, tumour size, tumour stage, tumour grade, and tumour content. These analyses showed that elevated IMPRES scores were associated with molecular subtype and age, but not with other clinical or demographic factors (Supp. Table 5). Following up on this result, multiple linear regression analysis of immune scores against all available molecular data, adjusting for age and molecular subtype, showed that mutational signature 3, tumour heterogeneity (PyClone clusters), and predicted neoantigen binding to be independently associated with IMPRES scores (Supp. Table 6), although the overall predictive ability of the model was weak (adjusted R^2^ = 0.13).

Finally, to determine the biological pathways that are associated with higher immune scores, we conducted gene set enrichment analysis (GSEA) comparing tumours in the top quartile of IMPRES scores to tumours in the bottom quartile. We found that the systemic lupus erythematosus (SLE) pathway was the most enriched pathway for tumours in the top IMPRES score quartile (Supp. Table 7). Additionally, we also found enrichment in the cGAS-STING cytosolic DNA-sensing pathway, and decreased expression of the TGF-β signalling pathway in the top quartile, which is concordant with previous studies on the tumour immune microenvironment (Supp. Table 7)^36–38^. We confirmed these results by comparing IMPRES scores for our cohort to GSVA scores using the KEGG gene sets, and found a strong correlation between the two scores for all three pathways (SFigure 25).

### Clinical impact of genomic profiles of Asian breast cancer

To determine the potential clinical impact of the differences in molecular profiles of breast cancers in Asian women, we determined the impact of the observations on overall survival. As expected, we found that ER-patients had poorer survival compared to ER+ patients, patients with IntClust 10 had poorer survival relative to other IntClusts, and patients with basal tumours by PAM50 had poorer survival relative to other molecular subtypes (Figure 5A). In ER+ patients alone, *TP53* mutations were associated with poorer survival (Figure 5B). Patients with high IMPRES scores had significantly better survival in both unadjusted (Figure 5B) and adjusted (SFigure 26) models, and the effect was stronger in ER+ patients (Figure 5B). Interestingly, ER+ patients with both a low IMPRES score and *TP53* mutation appear to have markedly poorer survival than other patients (Figure 5B), although the sample size was small (n=8). Overall, these data suggest that the differences in molecular profiles of Asian breast cancer patients could have important clinical implications, particularly in patients with ER+ tumours.

**Figure 5:**
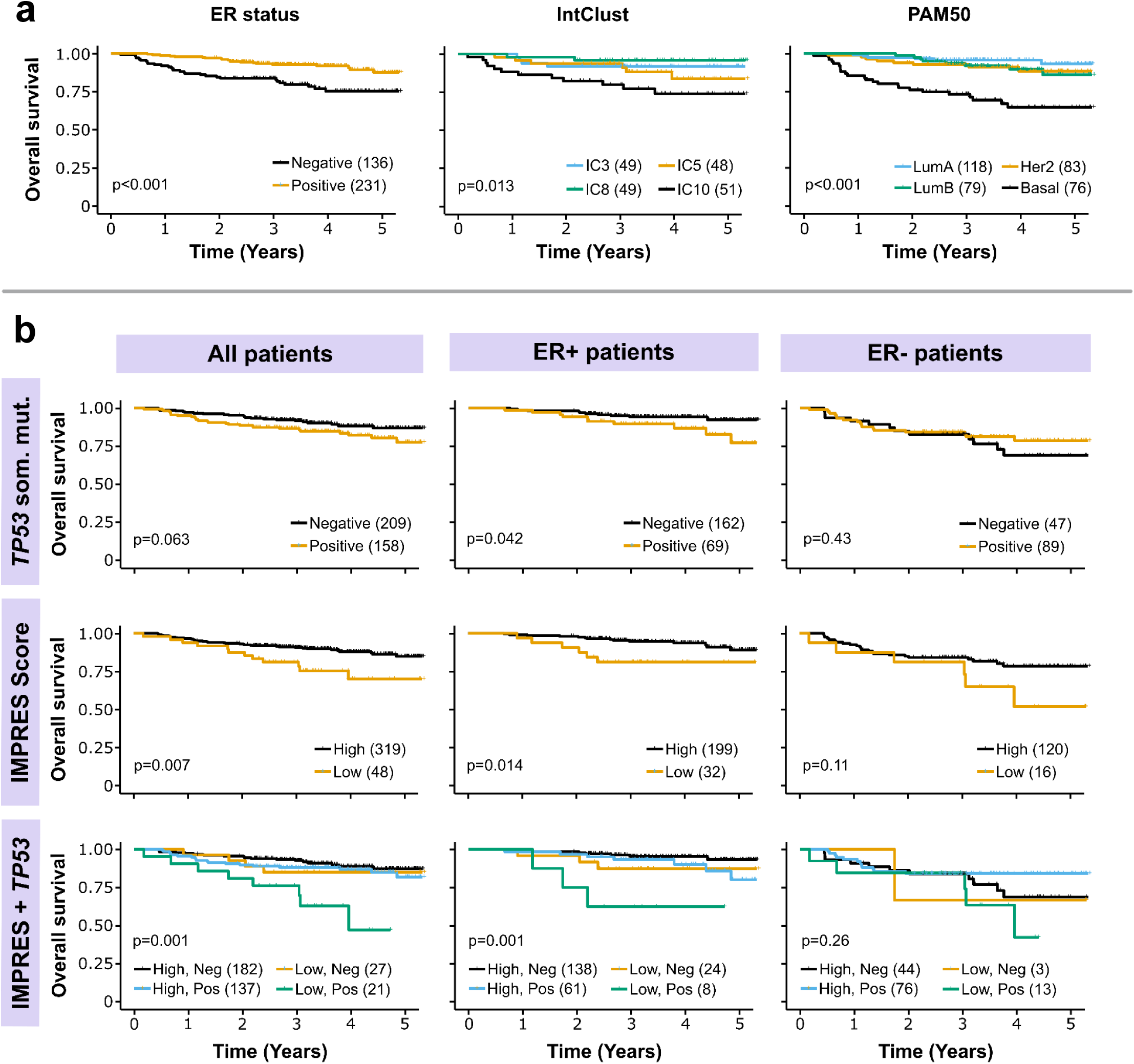
Survival analyses of MyBrCa patients. **a)** 5-year overall survival of MyBrCa patients stratified according to ER status (left), the four most common Integrative Clusters (middle), or PAM50 subtype (right). **b)** 5-year overall survival of MyBrCa patients stratified according to presence or absence of *TP53* somatic mutations (top), tumour IMPRES score (“High” cutoff = 9; middle), or a combined analysis of IMPRES score and *TP53* somatic mutations (bottom). Results are shown for all patients (left), as well as for only ER+ patients (middle), or ER-patients (right). For all figures, only patients with more than two years of follow-up data were included. Brackets in the figure legends indicate sample size for each group, and p-values shown are for unadjusted log-rank tests. Crosses indicate patient dropouts.

## DISCUSSION

To our knowledge, this is the largest and most extensive molecular profiling study on breast tumours arising in Asian women. Our study complements previous genomics studies of Asian breast tumours from China and South Korea^9,19,21^, and more importantly, enables more comprehensive comparisons with breast tumours arising in women of European descent^5,10,21^.

Comparisons conducted in this study revealed a higher prevalence of the Her2 molecular subtypes, higher prevalence of *TP53* mutations in ER+ disease and higher overall immune signatures in Asian breast cancer, all of which could impact clinical outcomes.

The higher prevalence of the Her2 subtype in Asian breast cancer is consistent with immunohistochemistry-based epidemiological studies among Asians and Asian Americans^39–42^. It is becoming increasingly apparent that the lifestyle and genetic risk factors may have overlapping but sometimes distinct effects in different subtypes of breast cancer. In fact, a large study of 28,095 breast cancer patients has suggested that parity may be associated with early onset Her2-positive breast cancers^43^. Incidentally, the MyBrCa cohort^20^ has a higher median parity relative to the Caucasian cohort sequenced by Nik Zainal et al. (2016)^21^. Further work on the basis for population differences in risk is warranted to understand implications for prevention.

The higher prevalence of *TP53* mutations in ER+ breast cancer in Asians is consistent with previous smaller studies in other Asian populations^19,29^. However, unlike liver cancer where *TP53* mutations may be linked to population-specific exposures to carcinogens^44^, we found no difference in the type or location of mutations nor in mutational signatures in Asian and Caucasian breast tumours, suggesting that the source of mutations may be similar in both populations. Regardless of the cause of the mutations, *TP53* mutations have been suggested to be prognostic, but with opposite effects in different subtypes of breast cancer^45,46^, and in ER+/HER2-disease, *TP53* mutations appear to be associated with higher risk to recurrence and poorer prognosis^47–49^. Notably, expression of *TP53* mutations may have different effects in different populations^50^, highlighting the need for further clinical studies in diverse populations.

Finally, we observed an elevated immune score, driven by higher cytotoxic TILs, in Asian breast tumours relative to the Caucasian tumours and in our cohort, elevated immune score is associated with better overall survival. Our results are consistent with recent microarray-based analyses in Asian breast cancer patients^51^. Intriguingly, ethnic-specific differences in the tumour microenvironment, such as hypoxia have been reported and appeared to be associated with higher *TP53* and lower *PIK3CA* mutations in Asians^52^. We found that the elevated immune scores are associated with the SLE, cGAS-STING cytosolic DNA-sensing, and TGF-β signalling pathways, HR deficiency, neoantigen prediction, and tumour heterogeneity. However, these associations only account for a small proportion of the population variance in immune scores, highlighting the need for further studies in this area.

In summary, this study may have important implications on our understanding of breast cancers arising in different populations. It has been reported that Asian-American breast cancer patients have better survival than other racial/ethnic groups^53,54^, and the molecular basis of breast cancers in Asian women reported in this study highlights some of the possible explanations for these population-specific differences in outcome.

## Data Availability

Access to controlled patient data requires the approval of the Data Access Committee. Requests can be submitted to.

## Acknowledgements

This project was funded by a research grant from the Newton-Ungku Omar Fund (MRC Ref: MR/P012442/1). Cancer Research Malaysia also receives charitable funding from the Scientex Foundation, Estée Lauder Companies, Yayasan Petronas, and Yayasan Sime Darby which contributed to the funding of this study. SFC, OMR, and CC also receive funding from Cancer Research UK. The authors would like to thank Dr. Tan Min Min for help with data curation, nurses and staff who helped with sample collection, as well as Dr Saira Bahnu Mohamed Yousoof and all staff at the Subang Jaya Medical Centre Tissue Diagnostics laboratory for assistance with histopathological sample retrieval and processing. The authors would also like to acknowledge the contribution of the Core Genomics Facility at the CRUK Cambridge Institute, where the sequencing work was conducted.

## Author Contributions

JWP and MMAZ co-led the data analysis and wrote the manuscript. PSN, MYM, SNH, and CHY contributed to sample collection and processing and data collection, while BS, OMR, and SFC generated and collected data. SJS interpreted results and guided the data analysis. PR provided histopathology expertise, and together with CHY collected clinical data. OMR, CC, SFC, and SHT designed experiments, interpreted results, and drafted the manuscript. The project was directed and co-supervised by CC, SFC and SHT, and were responsible for final editing.

## Methods

### Biospecimen Collection and Pathological Assessment

661 tissue samples were collected from breast cancer patients recruited as part of the MyBrCa study at the Subang Jaya Medical Centre (SJMC), Subang Jaya, Malaysia. The project was reviewed and approved by the Independent Ethics Committee, Ramsay Sime Darby Health Care (reference no: 201109.4 and 201208.1), and written informed consent was given by each individual patient. Matching blood samples were obtained prior to surgery, while tumour samples were sectioned from the primary tumour during surgery and were immediately frozen and stored in liquid nitrogen.

Patients were excluded from this study for the following criteria: No corresponding tumour samples (n=5), no corresponding germline samples (n=5) and those who withdrew consent (n=12). Tumour samples were further excluded after clinicopathological review if they were found to be from rare histological subtypes and other breast diseases (n=5). Tumour samples were then sectioned for DNA and RNA extraction, and the top and bottom sections were stained with haematoxylin and eosin (H&E) and reviewed for tumour content. Tumour samples with an average tumour content of <30% (n=50) and those with insufficient DNA (n=8) were excluded from the study.

A small minority of patients that were included in this study (n=26) received neoadjuvant chemotherapy prior to tissue collection. Tissue samples also included 4 cases of bilateral breast cancer and 1 case of recurrence.

### Sample Selection and Genomic Material Extraction

DNA from blood samples was extracted using the Maxwell 16 Blood DNA Purification Kit with the Maxwell 16 Instrument, according to standard protocol. DNA from tumour samples was extracted with the QIAGEN DNeasy Blood and Tissue Kit according to standard protocol, followed by quantitation using the Qubit HS DNA Assay kit and Qubit 2.0 fluorometer (Life Technologies Inc). RNA from tumour samples was extracted using the QIAGEN miRNeasy Mini Kit with a QIAcube, according to standard protocol. Total RNA was quantitated using Nanodrop 2000 Spectrophotometer and RNA integrity was confirmed using Agilent 2100 Bioanalyzer. For DNA samples, only samples with a concentration above 20.0 ng/µL were included for sequencing. For RNA samples, only samples with a concentration of at least 20.0 ng/µL and a RIN number above 7 were included for sequencing.

### Next-Generation Sequencing Data Generation

#### DNA Sequencing Libraries

DNA libraries were generated from 50ng of genomic DNA using the Nextera Rapid Capture Exome kit (Illumina, San Diego, USA) as per manufacturer’s instructions. Prior to exome capture, 4nM pools of DNA libraries (n=48) was subjected to single end 50 shallow whole genome sequencing. Exome capture was performed in pools of 3 and subjected to paired end 75 sequencing on a HiSEQ4000 platform (Illumina, San Diego, USA).

#### RNA Sequencing Libraries

RNA libraries were prepared from 550ng of total RNA using the TruSeq Stranded Total RNA HT kit with Ribo-Zero Gold (Illumina, San Diego, USA) as per manufacturer’s instructions and subjected to paired end 75 sequencing on a HiSEQ4000 platform (Illumina, San Diego, USA).

### RNA Sequencing Analysis

#### Alignment and Quality Assessment

RNAseq reads were mapped to the hs37d5 human genome and the ENSEMBL GRCh37 release 87 human transcriptome using the STAR aligner (v. 2.5.3a)^55^. Samples with less than 15 million mapped fragments were excluded from further RNA-seq analyses. Gene-level aligned fragment counts were generated using featureCounts (v. 1.5.3), while gene-level expression in transcripts-per-million (TPM) was calculated using RSEM^56^. Genes with an average count of less than 10 or an average TPM score of less than 0.1 were filtered out and excluded from further analyses. Variant calling from RNA-seq data for sample ID reconfirmation and downstream analyses was also conducted using the GATK Best Practices workflow for RNA-seq– in brief, STAR-mapped reads were processed to mark duplicates with Picard, followed by exon splitting & trimming and base recalibration using GATK, and lastly variants were called using HaplotypeCaller. Sample identities were verified as described below in the WES section.

#### Molecular Subtyping

Gene-level count matrices for the SD and TCGA cohort were transformed into log_2_ counts-per-million (logCPM) using the voom function from the limma (v. 3.34.9) R package. The transformed matrices were then was subtyped according to PAM50 and SCMgene designations using the Genefu package in R (v. 2.14.0). Additionally, each matrix was quantile-normalized and subtyped according to integrative clusters using the iC10 R package (v. 1.5). For the METABRIC cohort, the normalized microarray expression matrix from the discovery set was used for Genefu and ic10 subtyping without modification. For IntClust 4, we designated each sample as being ER+ or ER-by fitting a 2-component mixture model to the distribution of *ESR1* expression in TPM using an expectation-maximization algorithm as implemented in the mixtools (v. 1.1) package in R.

#### Unsupervised cluster analysis

K-means consensus clustering was conducted using gene-median centred TPM gene expression scores for the MyBrCa and TCGA Caucasian cohorts, using the ConsensusClusterPlus (v.1.46)^57^ package in R.

#### Profiling the Tumor-Immune Microenvironment

Overall immune cell infiltration in the bulk tumour samples was assessed from RNA-seq TPM gene expression scores using ESTIMATE (v. 1.0.13)^28^, as well as with GSVA (v. 1.26) using the combined immune cell gene sets from Bindea et al. 2013^58^. For each sample, we also scored the immune features predictive of checkpoint inhibitor immunotherapy using IMPRES scores^29^ (only 14 out of 15 of the predictive features were available in our datasets) as well as with GSVA using the Expanded IFN-gamma gene set from Ayers et al. 2017^59^. The relative abundance of specific immune cell populations was estimated from RNA-seq TPM scores with the CIBERSORT^30^ and TIMER^60^ web tools, as well as through GSVA with individual immune gene sets from Bindea et al. 2013^58^.

#### Multivariate Analyses of Clinical and Molecular Features

To assess the associations between various clinical and molecular features, we performed regression analyses using the base stats package in R (v. 3.5.1). For continuous variables such as IMPRES score, regular linear regression was applied using the “lm” function to estimate regression coefficients and statistical significance.

### Shallow Whole Genome Sequencing

#### Alignment and CNA Assessment

The sequenced reads were mapped to the hg19 reference genome using bwa-mem^61^, sorted using samtools^62^ and dedupped using picard (http://broadinstitute.github.io/picard). Mapped reads were analyzed using QDNAseq^63^ to obtain 100kb segmented copy number profiles using standard protocol and default parameters. Copy number aberrations were called using CGHcall (v 2.40) as implemented in the QDNASeq R package, which calls each segment as normal, copy-number gain, copy-number loss, amplification or deletion using a mixture model. ENSEMBL hg19 genes with HUGO names were mapped to the segmented copy number calls by their start positions to determine the copy number status for each gene in each sample.

### Whole Exome Sequencing Analysis

#### Alignment and Quality Assessment

The sequenced reads were trimmed for Illumina adapters using trim_galore! (https://github.com/FelixKrueger/TrimGalore) and mapped to the human genome b37 plus decoy genome using bwa-mem version 0.7.12^64^. The mapped reads were merged and sorted using samtools^62^, and dedupped using picard (http://broadinstitute.github.io/picard). Realignment and base quality score recalibration were performed using GATK3^65^.

Realignment around indels was done concurrently on the tumour and normal pairs. For bilateral cases, the two tumours and normal were realigned simultaneously. Sample identities were verified by determining the concordance of the genotypes using GATK3 HaplotypeCaller^65^. Briefly, the haplotypes at 2,000 to 5,000 dbSNP positions were called for all samples (RNA-seq and WES experiments) in an all-versus-all manner. True match will typically have upwards of 95% concordance between samples from the sample individuals. Samples with ambiguous identities were excluded from further analyses.

#### Data Download

We downloaded the TCGA^10^ and METABRIC^5^ data via the Genomic Data Commons Data Portal or cBioportal^66,67^. We additionally downloaded data for Nik-Zainal et al.^21^ (from ftp://ftp.sanger.ac.uk/pub/cancer/Nik-ZainalEtAl-560BreastGenomes) and Zhengyan Kan et al.^19^ (from https://www.nature.com/articles/s41467-018-04129-4). For Nik-Zainal WGS somatic mutation analyses, we delimited our analysis to only the Nextera Rapid Capture regions with additional flanking 100bp.

#### Short Nucleotide Variants Calling

We first constructed panel-of-normals by artefact detection mode of GATK3 Mutect2^65^, retaining only sites that are present in 2 or more normal samples. In all steps, we only concentrated on the Nextera Rapid Capture regions with additional flanking 100bp. For SNVs, we used positions that are called by Mutect2, and applied following filters: minimum 10 reads in tumour and 5 reads in normal samples, OxoG metric less than 0.8, variant allele frequency (VAF) 0.075 or more, p-value for Fisher’s exact test on the strandedness of the reads 0.05 or more, and S_AF_ more than 0.75. For positions that are present in 5 samples or more, we removed two positions that were not in COSMIC and in single tandem repeats. We also removed variants that have VAF at least 0.01 in gnomAD^68^, and considered only variants that are supported by at least 4 alternate reads, with at least 2 reads per strand. For indels, we also required the positions to be called by Strelka2^69^. We manually salvaged an indel in PALB2 gene in sample SD0028 which we have identified using PCR. We annotated the variants using Oncotator version 1.9.9.0^70^. We plotted mutation positions as lollipop plots using MutationMapper^66,67^.

#### Cancer Cell Fractions

Cancer cell fractions were calculated for nine common breast cancer driver genes using VAFs, copy number and tumour purity estimates obtained from WES, following the methodology described in Pereira et al. (2016)^7^. Copy number and tumour purity were generated for this analysis using the ASCAT R package (v. 2.5.2) on allele counts generated from WES bam files using alleleCounter (v. 4.0.1).

#### Driver Genes Analysis

We considered samples from WGS and WES experiments (MyBrCa, TCGA Asian, Korean, WGS Asian, TCGA Caucasian, WGS Caucasian) for driver gene analyses^22^. We considered genes that are mutated in at least 1% of samples in Asian ER-, Asian ER+, Caucasian ER+ or Caucasian ER-. We calculated oncogene score (ONC) as the percentage of missense mutation counts at the most recurrent amino acid positions over all mutations, and considered only genes with at least 5 recurrent mutations. We calculated tumour suppressor score (TSG score) as the percentage of inactivating mutation counts over all mutations, and considered only genes with at least 5 inactivating mutations. We selected genes with either ONC or TSG scores of more than 20, and queried the Integrative Onco Genomics database (www.intogen.org)^71^ for known cancer driver genes (STable 2 and 3). We sorted 40 genes according to the enrichment in Asian ER+ over Caucasian ER+ and plotted the prevalence and the ONC/TSG scores (SFig. 6).

#### Association Patterns of Driver Genes

We considered the top 15 mutated genes from driver genes analysis. We calculated odds ratio and determined the likelihood of co-occurrence or mutual exclusivity of missense or inactivating mutations using Fisher’s exact test. We displayed associations with p-value of less than 0.05 after Bonferroni correction and plotted the log odds ratio.

#### Mutational Signatures

We used deconstructSigs^72^ to determine the weights of previously-reported breast cancer mutational signatures (Signatures 1, 2, 13, 3, 8, 6, 15, 20, 26, 5, 17, 18 and 30) from COSMIC matrices (version 2, March 2015) in samples with at least 15 sSNVs. We determined the proportions of mutational signatures in the different datasets by combining the variants from all samples, and used the function plotSignatures^72^ to plot the mutational spectra. To determine the difference in mutational signature weights between the top and bottom immune quartile of MyBrCa samples, we ranked the samples by their ESTIMATE scores, and performed 1-sided rank sum Wilcoxon’s test on the two categories.

#### Germline Mutation Analysis

Carriers of deleterious germline mutations in *BRCA1, BRCA2, TP53, PALB2, ATM*, and *CHEK2* were identified from targeted sequencing of the MyBrCa cohort (BRIDGES study, in review), which were confirmed with Sanger sequencing. Comparable data from TCGA was taken from Huang et al. (2018)^24^.

#### Correlation between Immune Scores and Mutational Signatures

We calculated Spearman’s correlation co-efficient between the mutational signatures and immune scores and plotted the correlations as heatmaps.

#### Differential Expression and Pathway Analyses

Gene-level count matrices were normalized using the “Trimmed Mean of *M*-values” (TMM) method implemented in the edgeR (v. 3.20.9) R package. The count matrices were then transformed into logCPM using the voom function from the limma package in R. Differentially expressed genes were determined by empirical Bayes moderation of the standard errors towards a common value from a linear model fit of the transformed count matrices as implemented in the limma package, with the threshold for differential expression set as FDR <0.001 and absolute log fold-change >0.2. For pathway analysis, DE genes were further filtered for FDR < 1e-50, and ranked according to absolute log fold-change. The top 1000 ranked genes were queried in Reactome (www.reactome.org) to determine the top enriched pathways. Further pathway analyses were also conducted using GSEA (v. 3.0)^73^ on quantile-normalized gene-level count data. GSEA was run for 1000 permutations with the v. 6.2 Hallmark and KEGG gene sets from MSigDB.

#### Validation of Immune Scores with Immuno-histochemistry

FFPE blocks for 207 patients with sequencing data were sectioned and stained for anti-CD3 (clone 2GV6, predilute; Ventana Medical Systems), anti-CD4 (clone SD35, predilute; Ventana Medical Systems), anti-CD8 (clone SD57, predilute; Ventana Medical Systems) and anti-PD-1 (clone SP263, predilute; Ventana Medical Systems) using an automated immunostainer (Ventana BenchMark ULTRA; Ventana Medical Systems, Tucson, AZ). Stained slides were digitized using an Aperio AT2 whole slide scanner. CD3, CD4, and CD8 staining was quantified using the Aperio Positive Pixel digital pathology tool and PDL-1 expression was determined using the Combined Positive Score system. The data were further verified by a pathologist (PR).

#### Quantification of Tumour Heterogeneity

Tumour heterogeneity was determined using PyClone (v 0.13.1)^33^ with default options to estimate the number of subclonal clusters within each tumour sample. Allele counts used for the PyClone input were extracted from the GATK output MAF files, while copy number input data was generated by ASCAT (v. 2.5.2) from WES allele counts generated by alleleCounter (v 4.0.1). Tumour heterogeneity was also separately quantified for each sample in the MyBrCa and TCGA cohorts using the MATH method described in Pereira et al. (2016)^7^ from MAF files from WES.

#### Neoantigen Analysis

Sample HLAs were determined using Polysolver^74^ from tumor and normal DNA WES data. Only HLA alleles that were concordant in tumor and normal WES data were considered. Somatic mutations were annotated using VEP^75^. All possible neoantigen peptides (9- to 11-mers) encompassing the nonsynonymous mutations were predicted using a combination of NetMHCpan^76^ and NetMHC^77^ on pVAC-seq platform^78^. Only neoantigens with predicted binding of less than 500nM were considered.

#### Survival Analysis

Overall survival data was obtained for each patient by querying their names and identity card numbers against the Malaysian National Registry records of deaths. Patients that did not return any matches against the database were assumed to still be alive, and *vice versa*. Length of survival was defined as the period of time from the date when patients were recruited into the study until the date of death as recorded by the Malaysian National Registry for patients who have passed away, or until the date when the Malaysian National Registry was last queried for patients assumed to still be alive. For all survival analyses in this study, only patients with at least two years of survival data were included (n = 367). Unadjusted Kaplan-Meier analyses and log rank tests were conducted using the survival package in R (v. 2.44) and plotted using the “ggtsurvplot” function from the survminer R package (v. 0.4.4). Cox proportional hazard models were built using the “coxph” function from the survival package and plotted using the “ggforest” function from survminer.

**SFigure 1:**
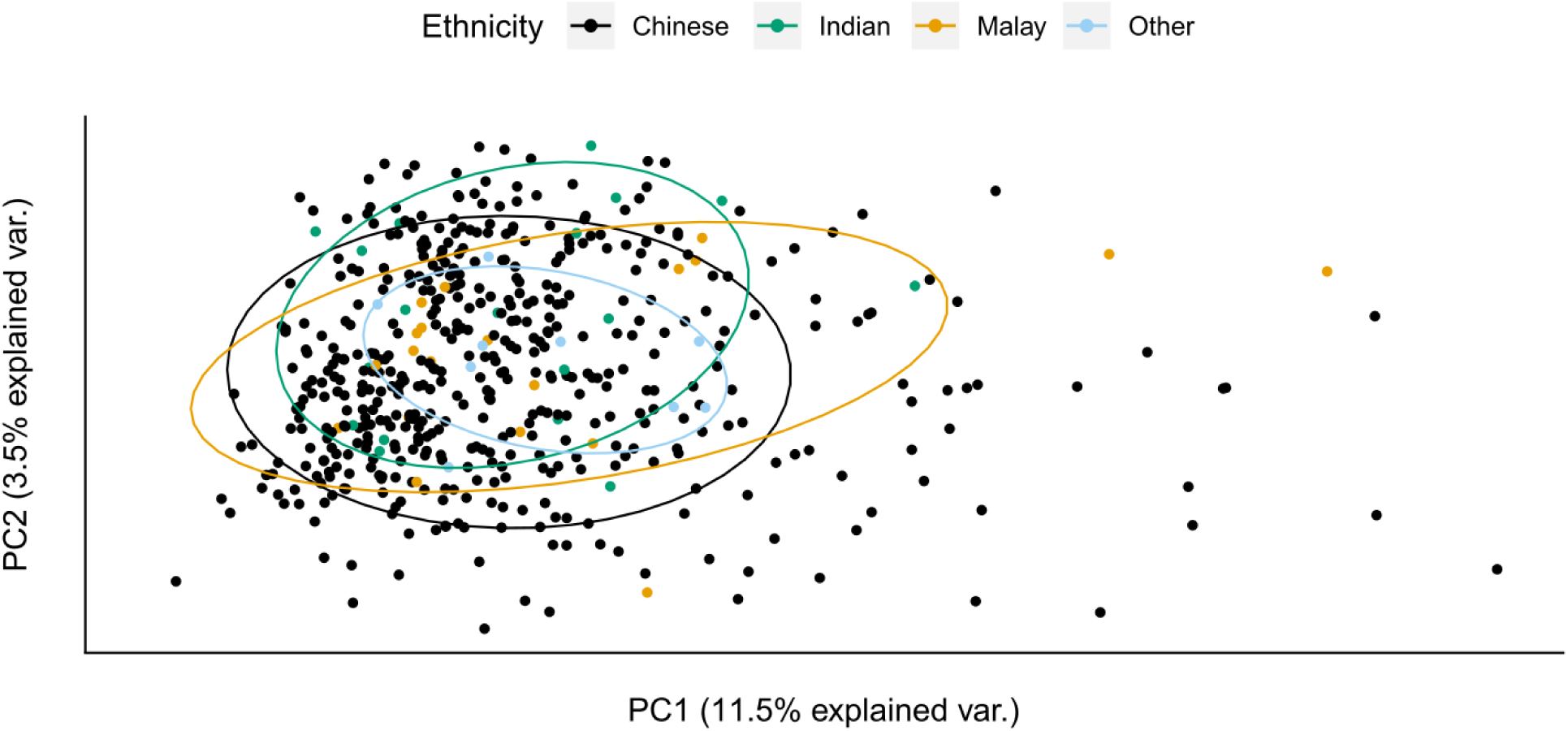
Ethnicity and gene expression in MyBrCa breast tumours. Principal component analysis of gene expression in MyBrCa tumours showing the overlap between different ethnicities (n: Chinese=498, Indian=19, Malay=26, Other=9).

**SFigure 2:**
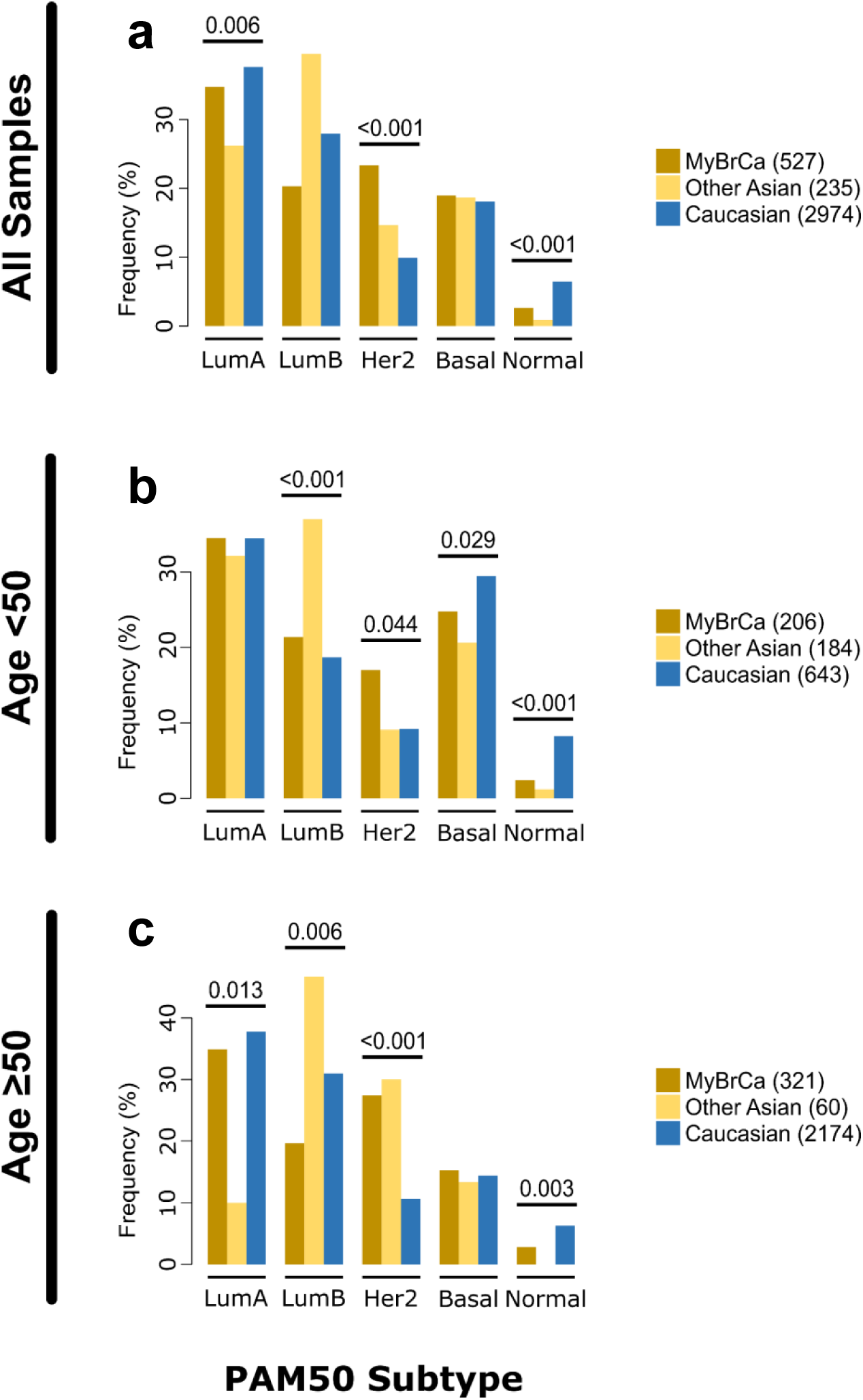
Molecular subtypes of Malaysian breast cancer. Comparison of PAM50 molecular subtype distribution across Malaysian (MyBrCa), other Asian (Korean, TCGA Asian), and Caucasian (TCGA Caucasian, METABRIC, Nik-Zainal 2016) cohorts. Comparisons were done using the full cohorts **(a)** as well as with only patients below **(b)** or above **(c)** the age of 50 as a rough proxy for menopausal status. Numbers above the bars are p-values denoting significant differences between Asians and Caucasians for that subtype, as determined by Pearson’s chi-square test. Numbers in the figure legend indicate sample size.

**SFigure 3:**
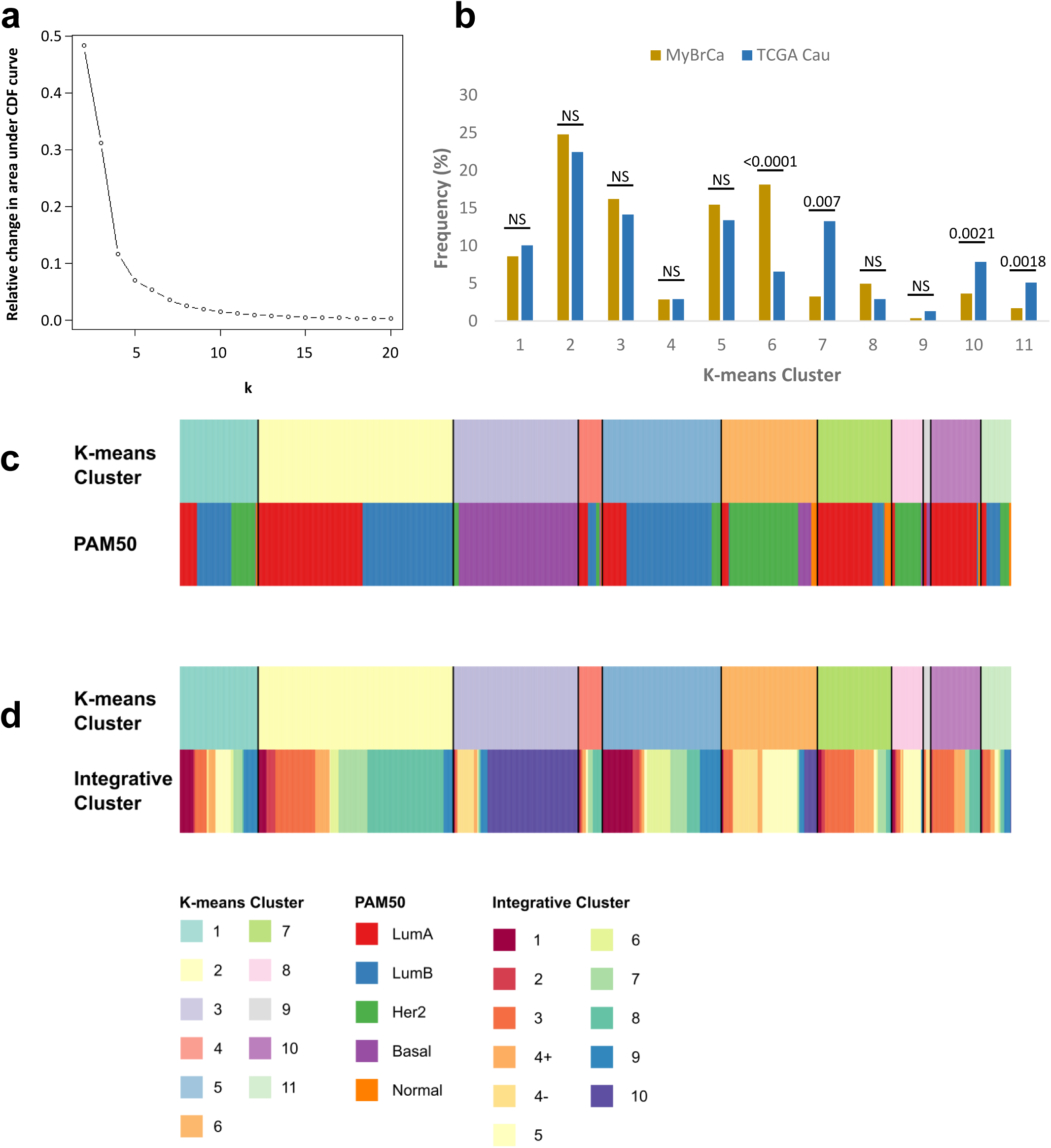
Unsupervised clustering of MyBrCa and TCGA Caucasian samples. K-means consensus clustering of MyBrCa and TCGA Caucasian gene expression data in order to determine the existence of exclusive Asian or Caucasian subtypes. **(a)** Plot of the relative change in area under the empirical cumulative distribution (CDF) of a consensus matrix, used to determine that the maximum number of meaningful clusters in our data is k=11. **(b)** Frequency of samples in each k-means cluster by cohort when k=11. **(c-d)** Comparison of the 11 k-means clusters to PAM50 (c) and Integrative Clusters (d).

**SFigure 4:**
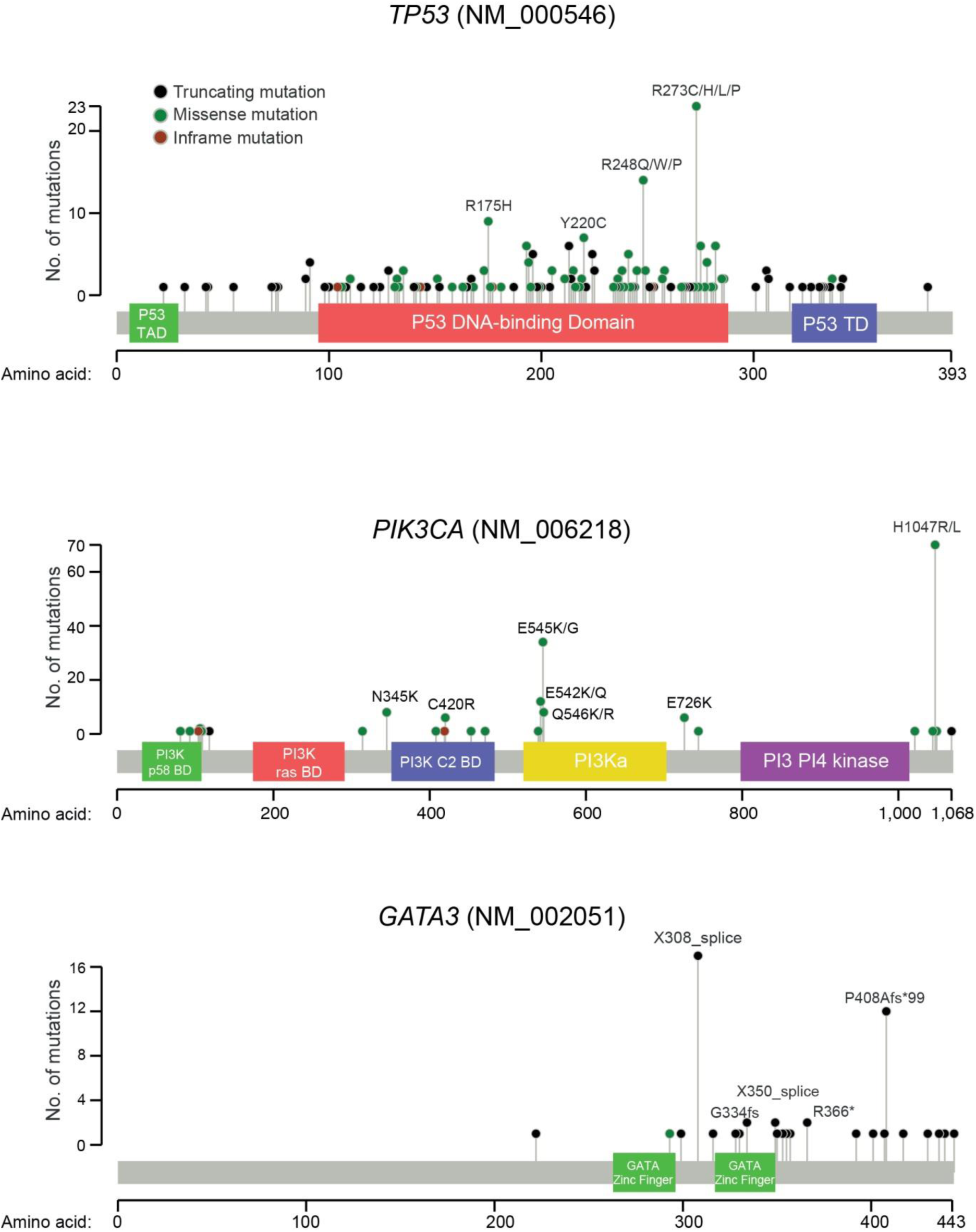
Somatic mutation positions of *TP53, PIK3CA* and *GATA3* in MyBrCa. Lollipop plots depicting positions of truncating, missense and inframe somatic mutations, as well as the most prominent positions in each gene.

**SFigure 5:**
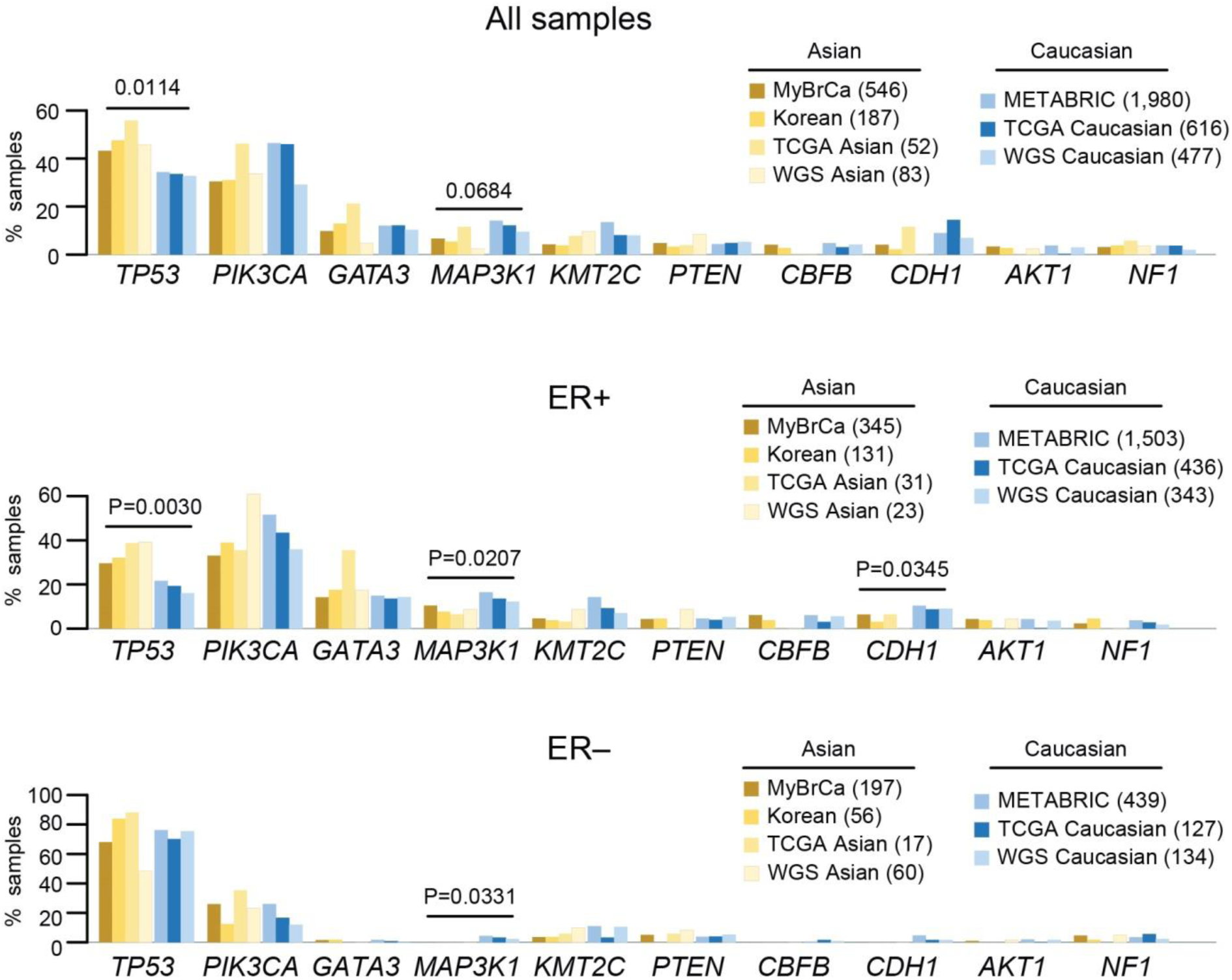
Mutational prevalence of main breast cancer genes in Asian and Caucasian breast tumours, separated by ER status. Comparison of mutational prevalence of main breast cancer genes in Asian (MyBrCa, Korean, TCGA Asian, WGS Asian) and Caucasian (METABRIC, TCGA Caucasian, WGS Caucasian) breast tumours, in all samples (top), or separated according to their ER status (middle and bottom). P-values from 2-sided Student’s *t*-test comparing Asian versus Caucasian samples.

**SFigure 6:**
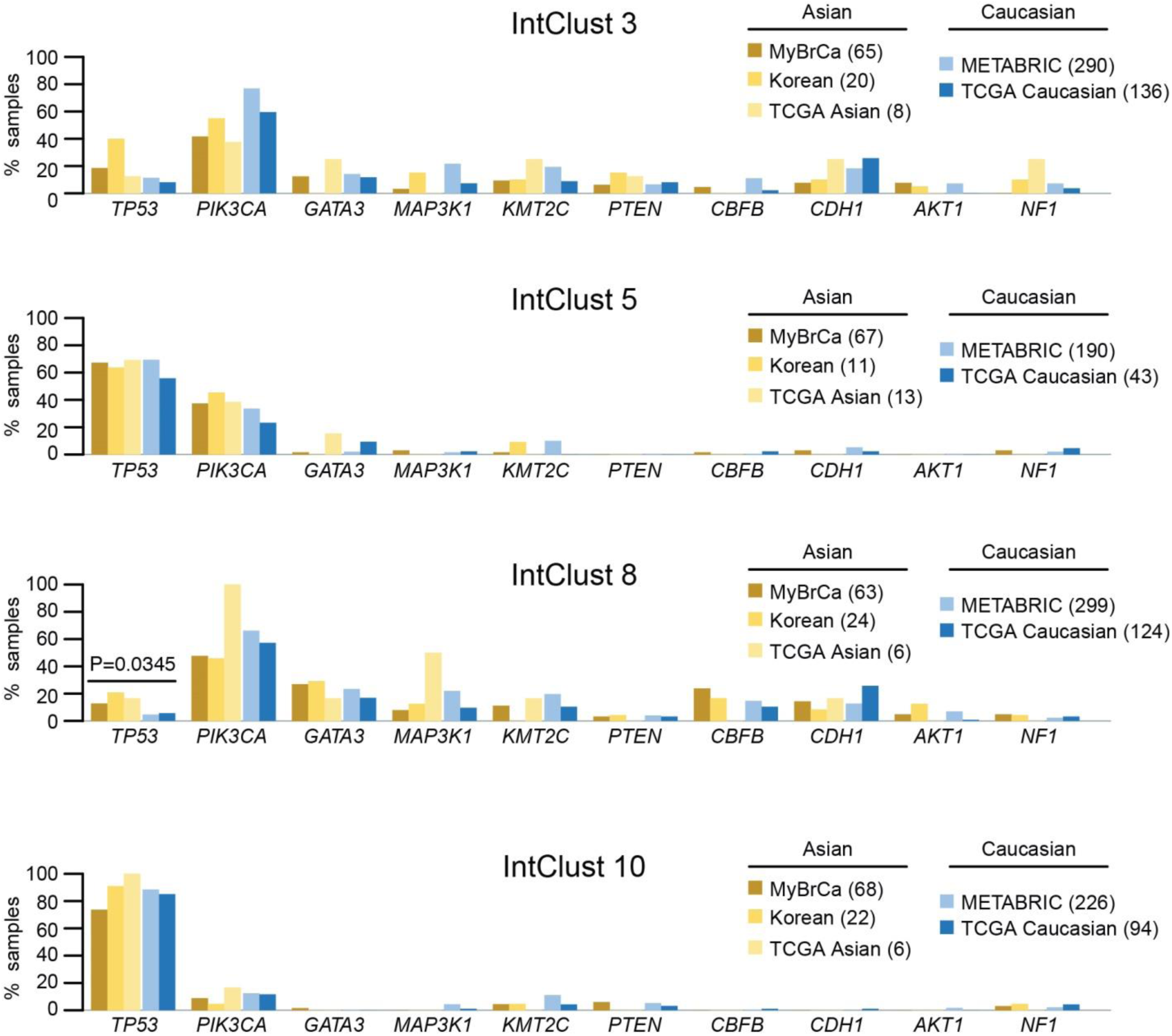
Mutational prevalence of main breast cancer genes in Asian and Caucasian breast tumours, separated by IntClust. Comparison of mutational prevalence of main breast cancer genes in Asian and Caucasian, limited to all samples from Integrative Clusters 3,5, 8 and 10 – these IntClusts are with the most samples overall to permit meaningful cross comparison. P-value from 2-sided Student’s *t*-test comparing Asian versus Caucasian samples.

**SFigure 7:**
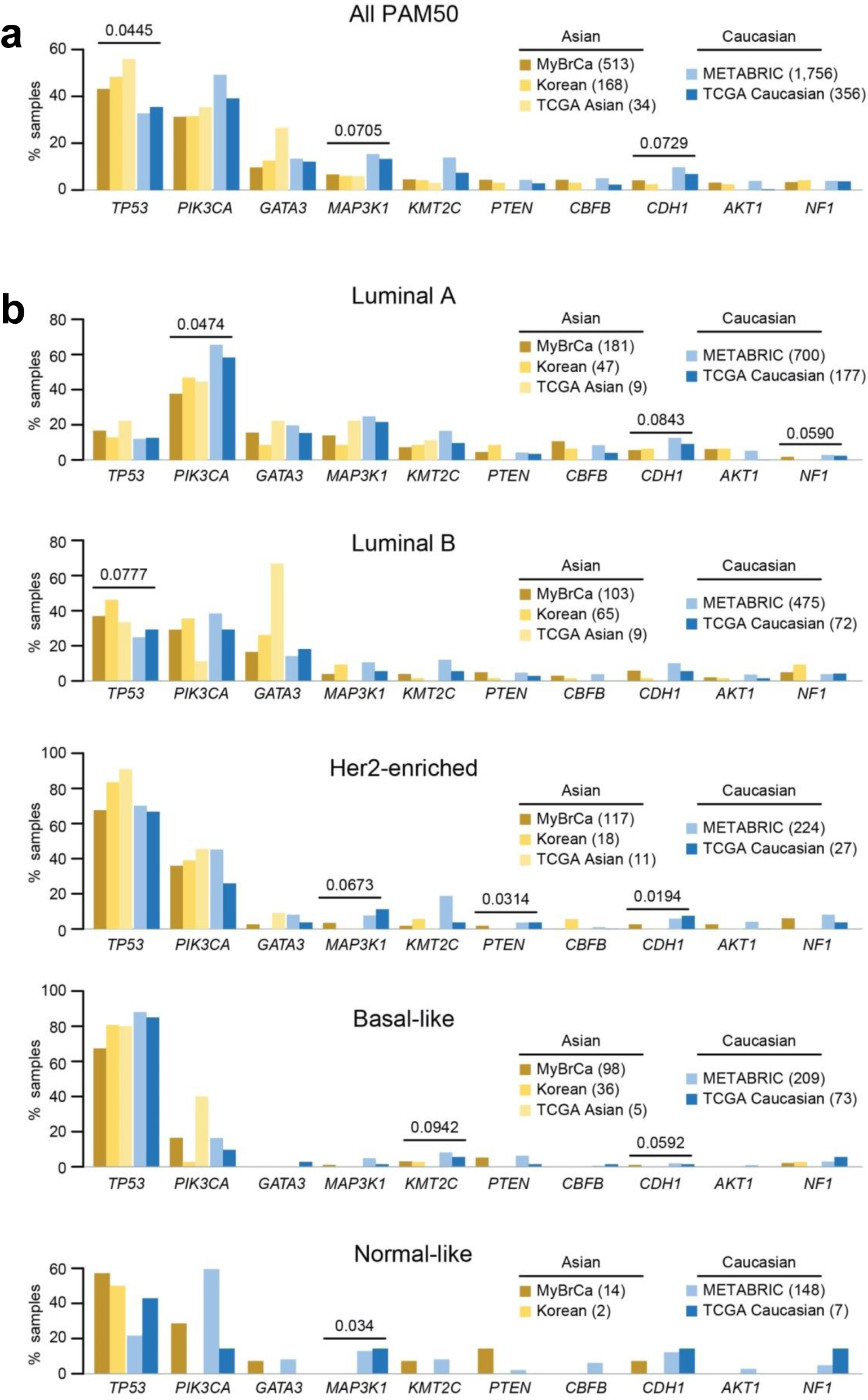
Mutational prevalence of main breast cancer genes in Asian and Caucasian breast tumours, separated by PAM50. Comparison of mutational prevalence of main breast cancer genes in Asian and Caucasian, limited to all samples with available PAM50 classification **(a)** and to PAM50 subtypes **(b)**. P-values from 2-sided Student’s *t*-test comparing Asian versus Caucasian samples.

**SFigure 8:**
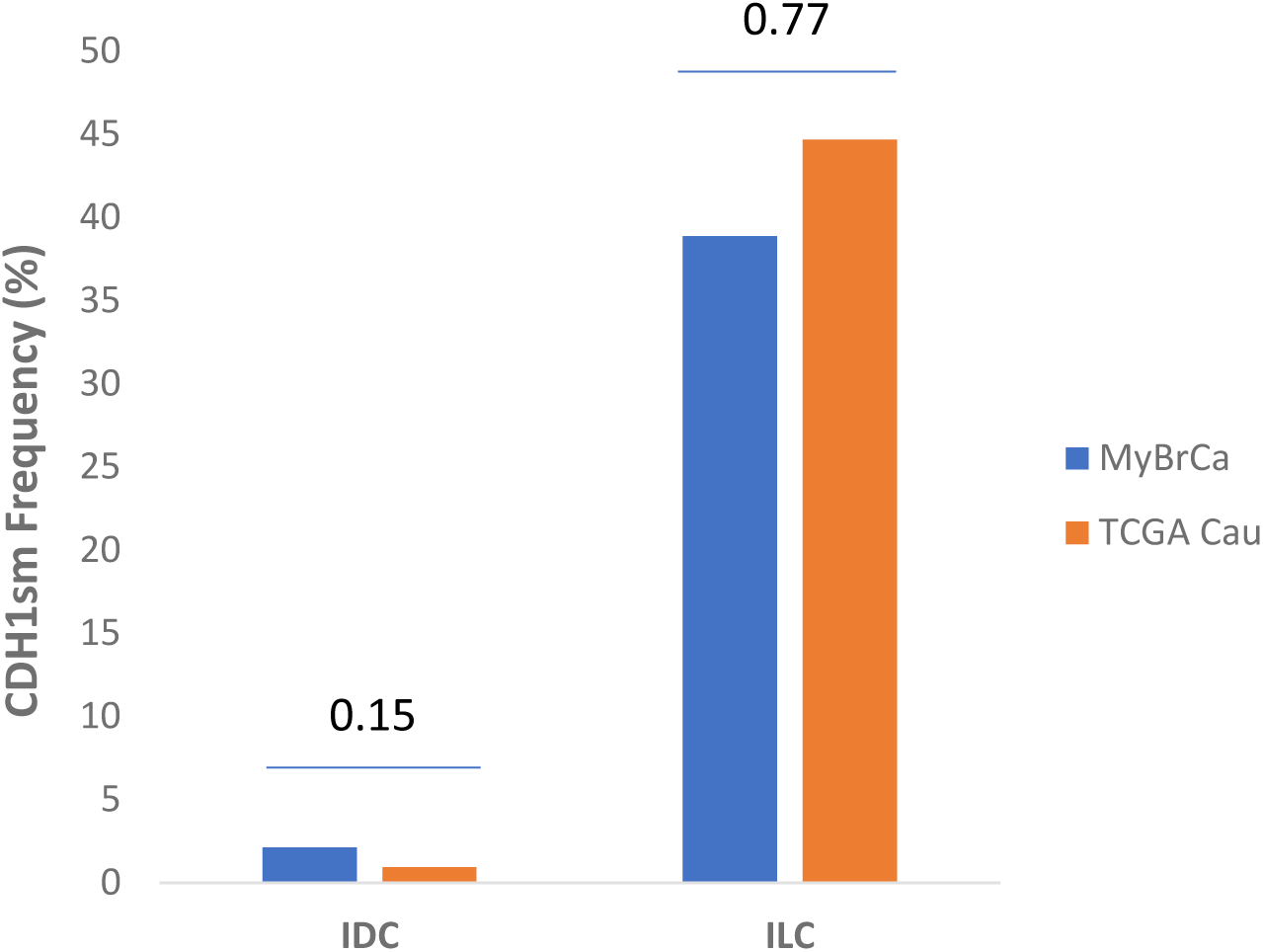
Prevalence of *CDH1* somatic mutations in different histological subtypes. Comparison of prevalence of somatic mutations in the *CDH1* gene in the MyBrCa and TCGA Caucasian cohorts, stratified by histological subtypes (IDC: Invasive ductal carcinoma; ILC: Invasive lobular carcinoma). P-values determined by chi-square test (n=533 and 574 for MyBrCa and TCGA Cau, respectively)

**SFigure 9:**
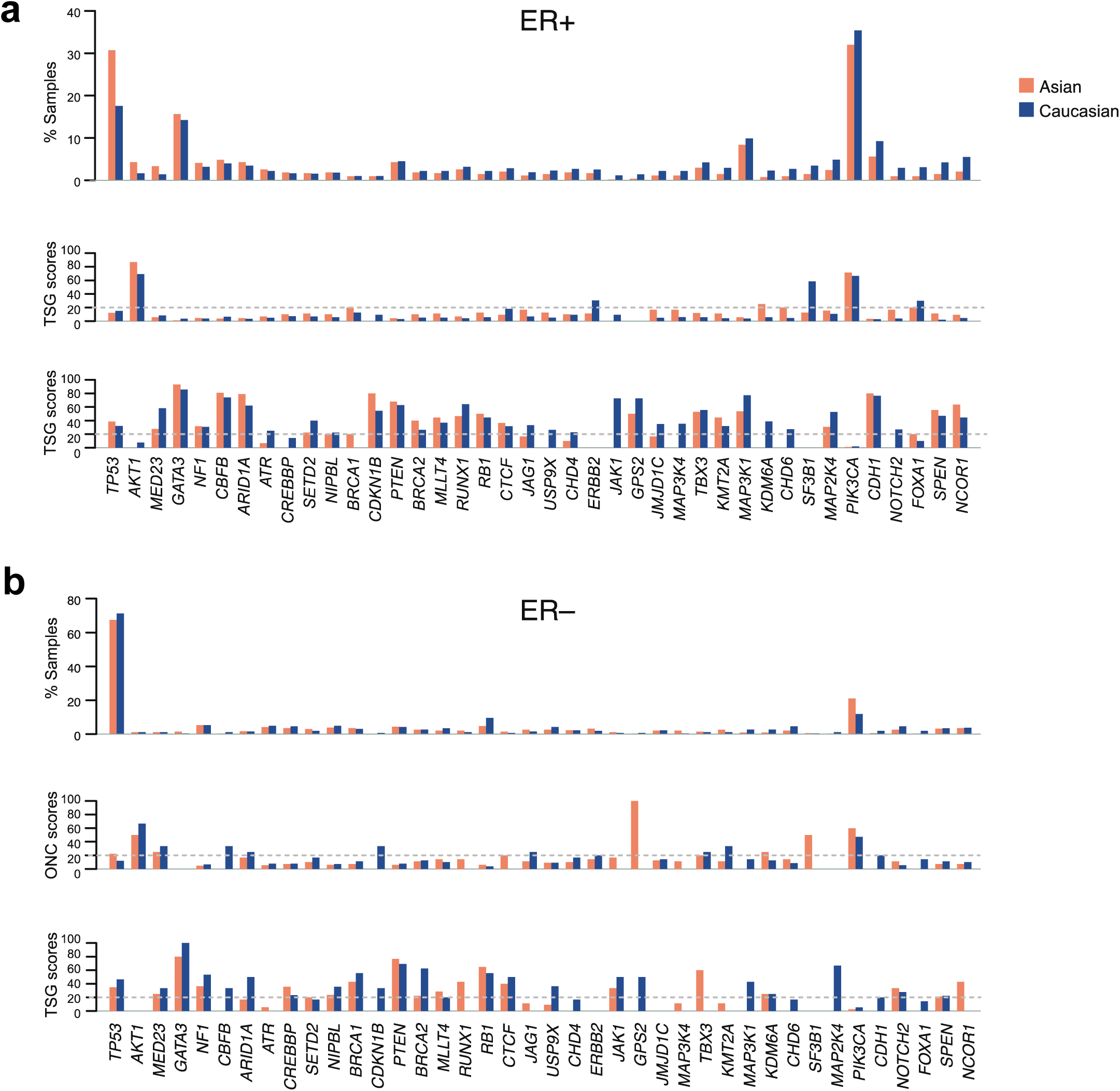
Driver genes in Asian and Caucasian breast tumours. Bar plots depict mutation rates of driver genes (top panels), as well as ONC (middle) and TSG (bottom) scores in Asian or Caucasian samples, separated into ER+ **(a)** or ER– **(b)**. Dotted grey lines are at 20 ONC/TSG scores.

**SFigure 10:**
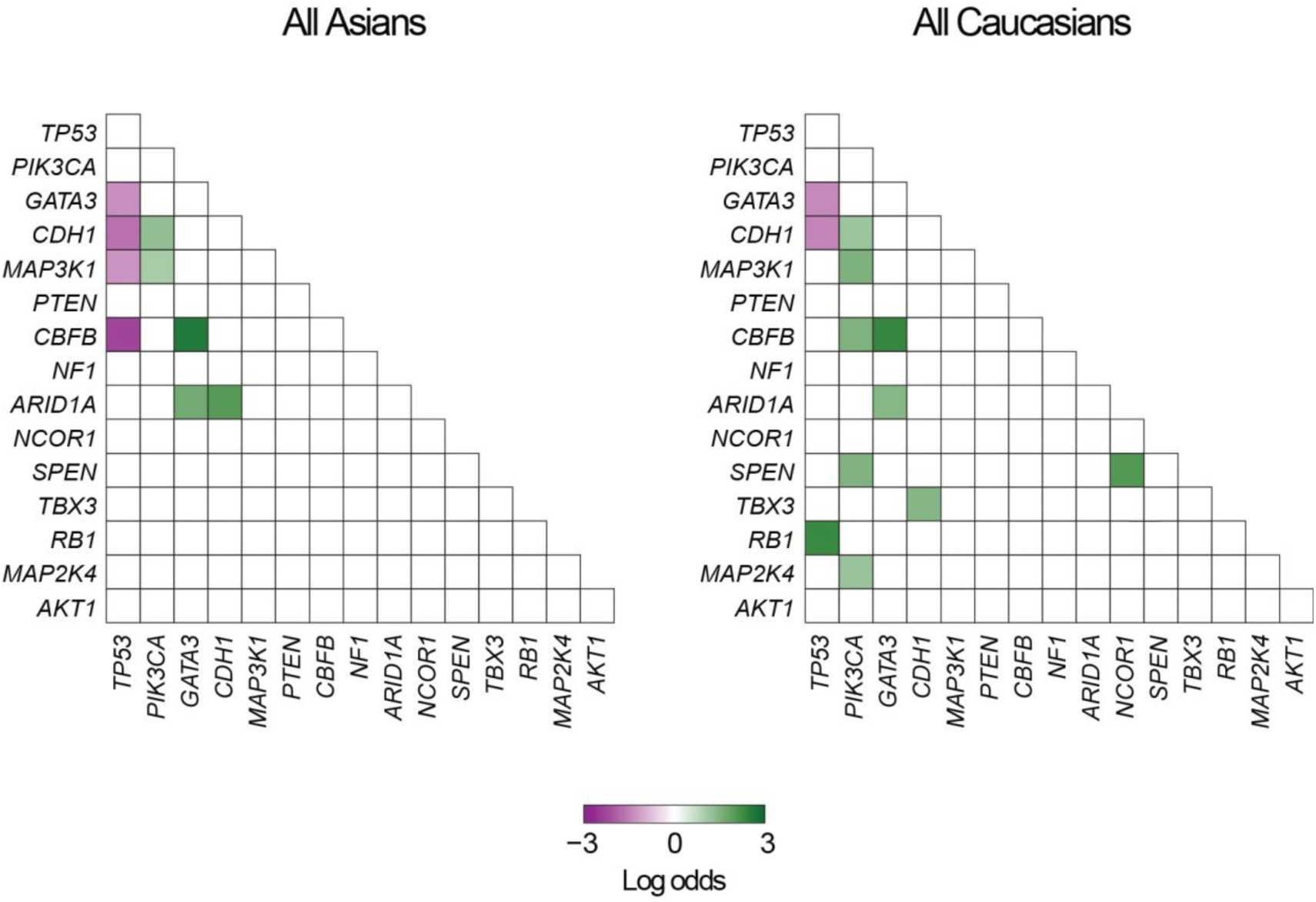
Association patterns of driver genes in Asian and Caucasian breast tumours. Heat maps depicting co-occurrence (green) or mutual exclusivity (purple) of somatic SNV or indel mutations in all Asian (left) and Caucasian (right) breast tumours.

**SFigure 11:**
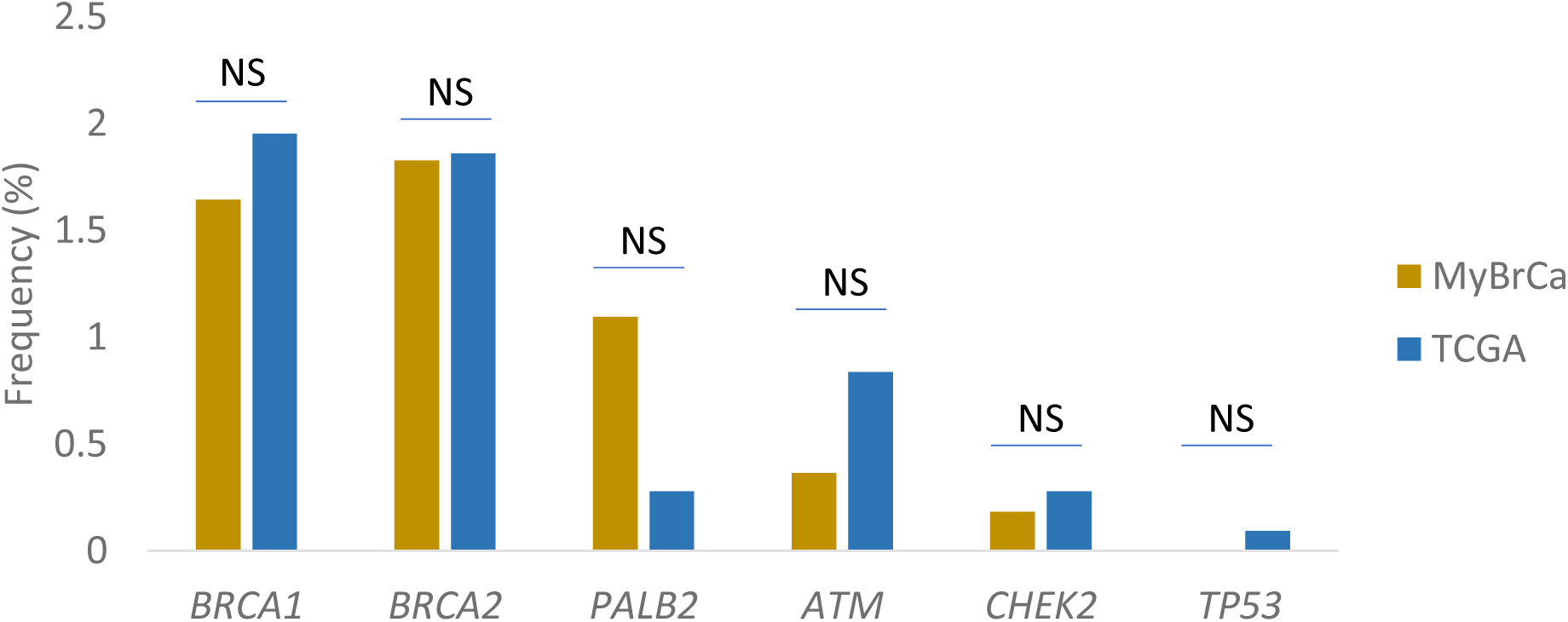
MyBrCa and TCGA breast tumours with germline pathogenic variants. Bar plots depict the frequency of tumours with germline pathogenic variants in 6 high to moderate breast cancer susceptibility genes. Significance determined by chi-square tests (NS: p<0.1). Germline pathogenic variants were defined as all frameshifts, stop-gain and splice site variants which result in loss of function as well as missense variants reported as pathogenic/ likely pathogenic in ClinVar.

**SFigure 12.**
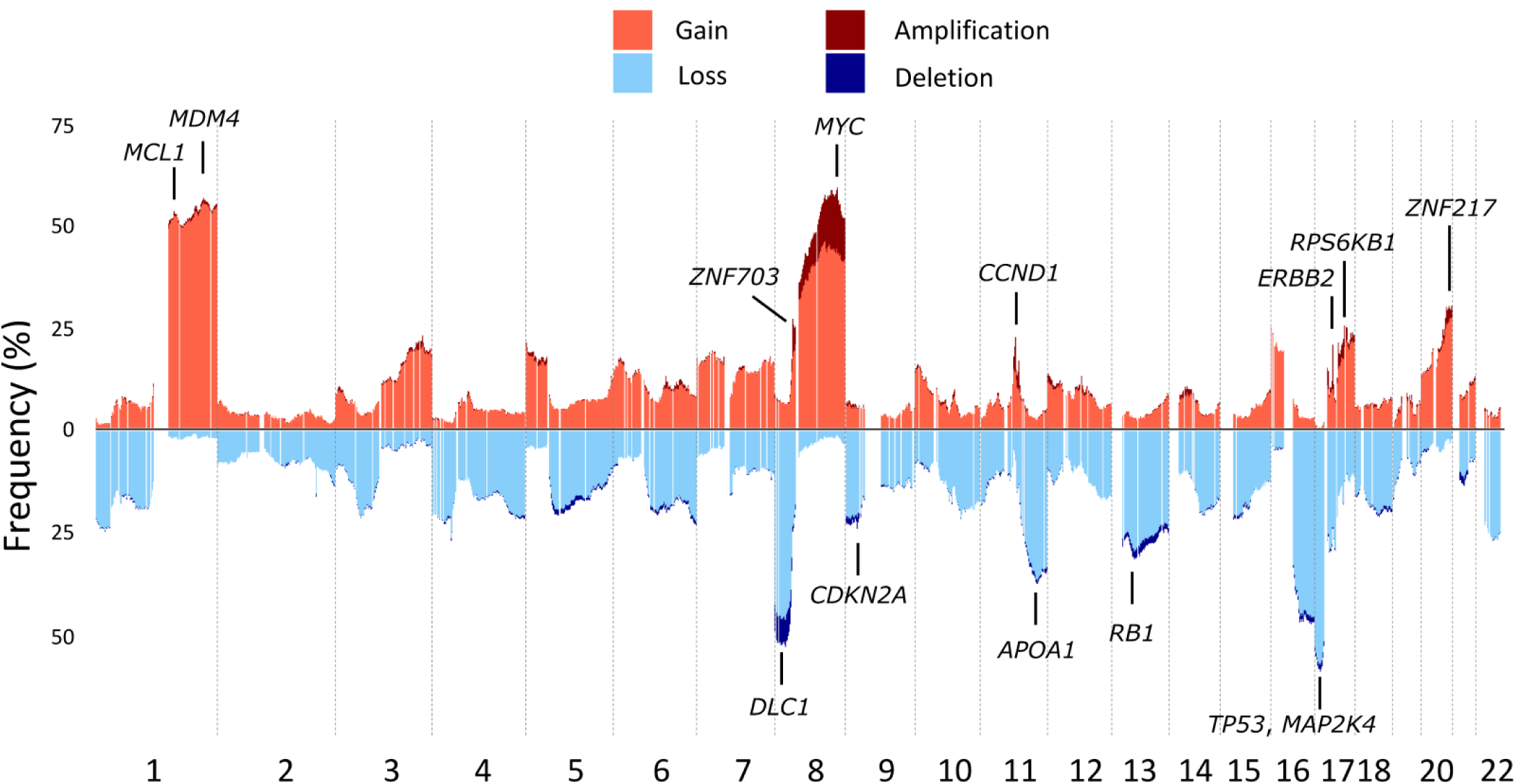
Copy number profile of MyBrCa tumours from shallow WGS. Frequency of copy-number amplifications, gain, loss, and deletions are based on 100kb-window segmented calls, as called by QDNASeq^66^.

**SFigure 13:**
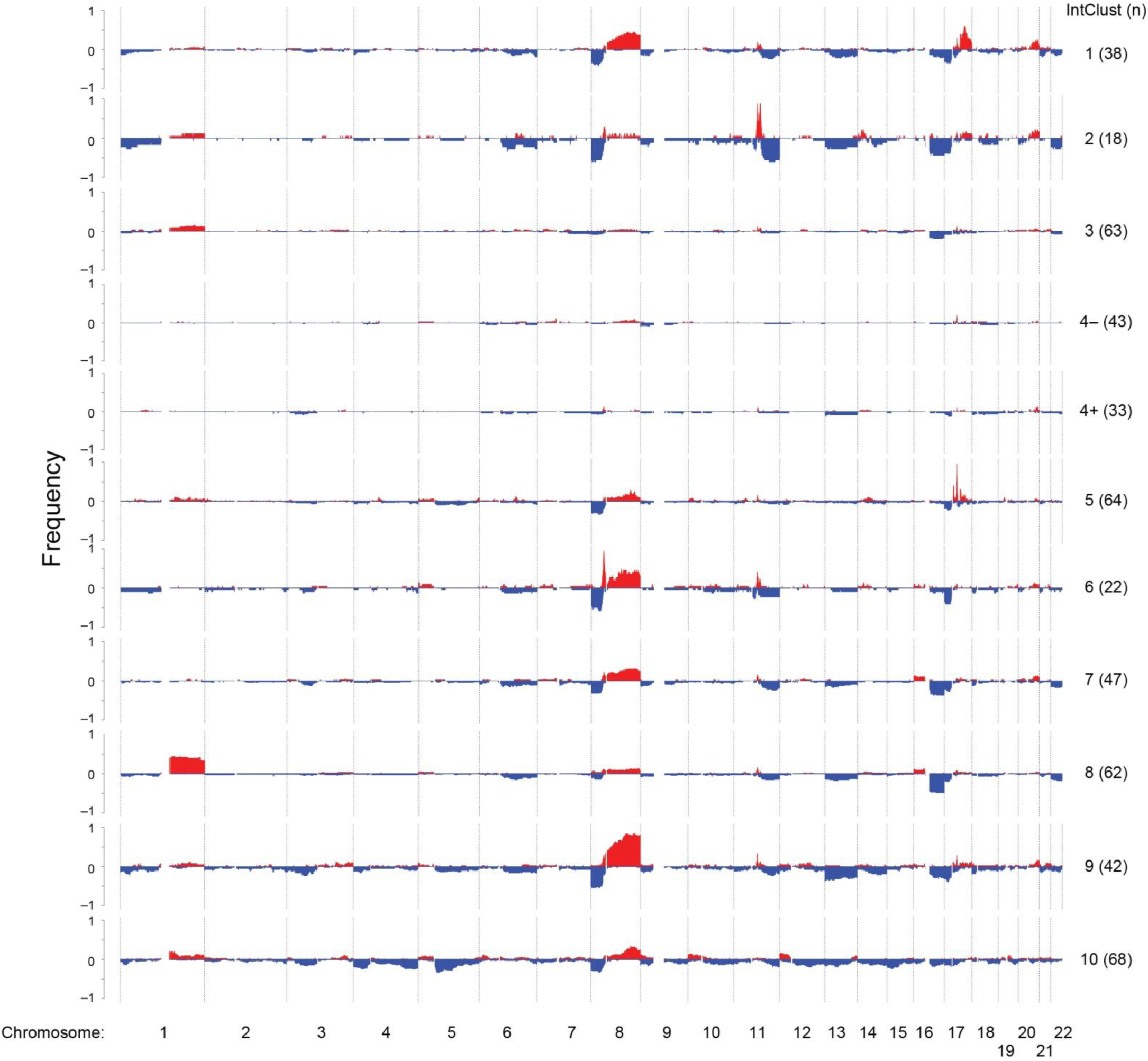
Copy number aberration (CNA) of MyBrCa samples. Copy number aberration plots depicting the frequencies of CNAs in each IntClust.

**SFigure 14:**
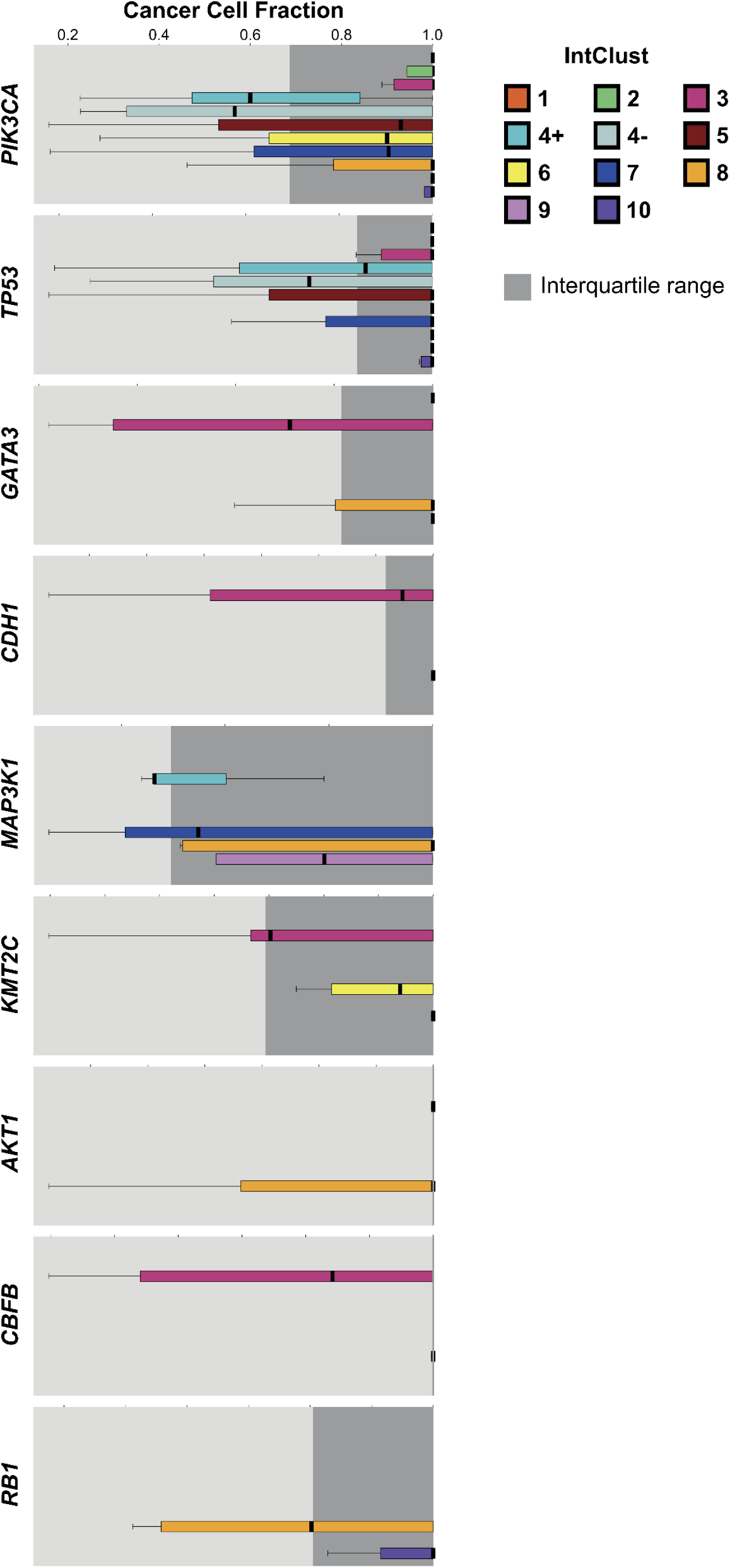
Cancer cell fraction of nine driver genes across Integrative Cluster subtypes in Malaysian breast tumours. Comparison of cancer cell fraction (CCF) – the proportion of cancer cells carrying a mutation for a specific gene – for nine common breast cancer driver genes across Integrative Cluster subtypes in the MyBrCa cohort. The interquartile range of CCF across all samples for each gene is shown in grey. Only subtypes with at least three samples in that category are shown.

**SFigure 15:**
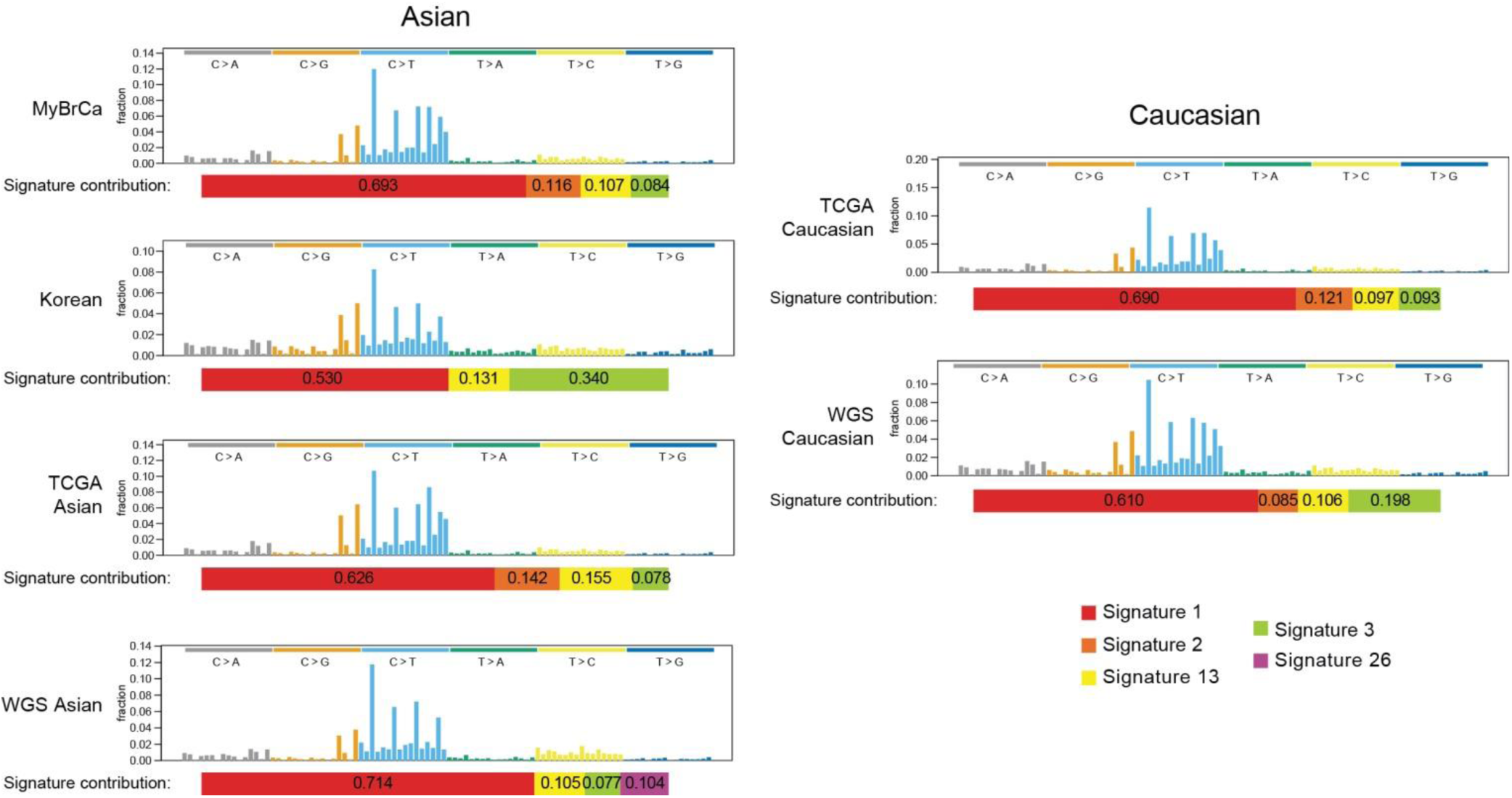
Mutational signatures of Asian and Caucasian breast tumours. Mutational spectra of Asian tumours (MyBrCa, Korean, TCGA Asian, WGS Asian) and Caucasian (TCGA Caucasian, WGS Caucasian) generated from all samples from each dataset. Also shown are the relative contributions of the mutational signatures in each data set (bottom bars).

**SFigure 16:**
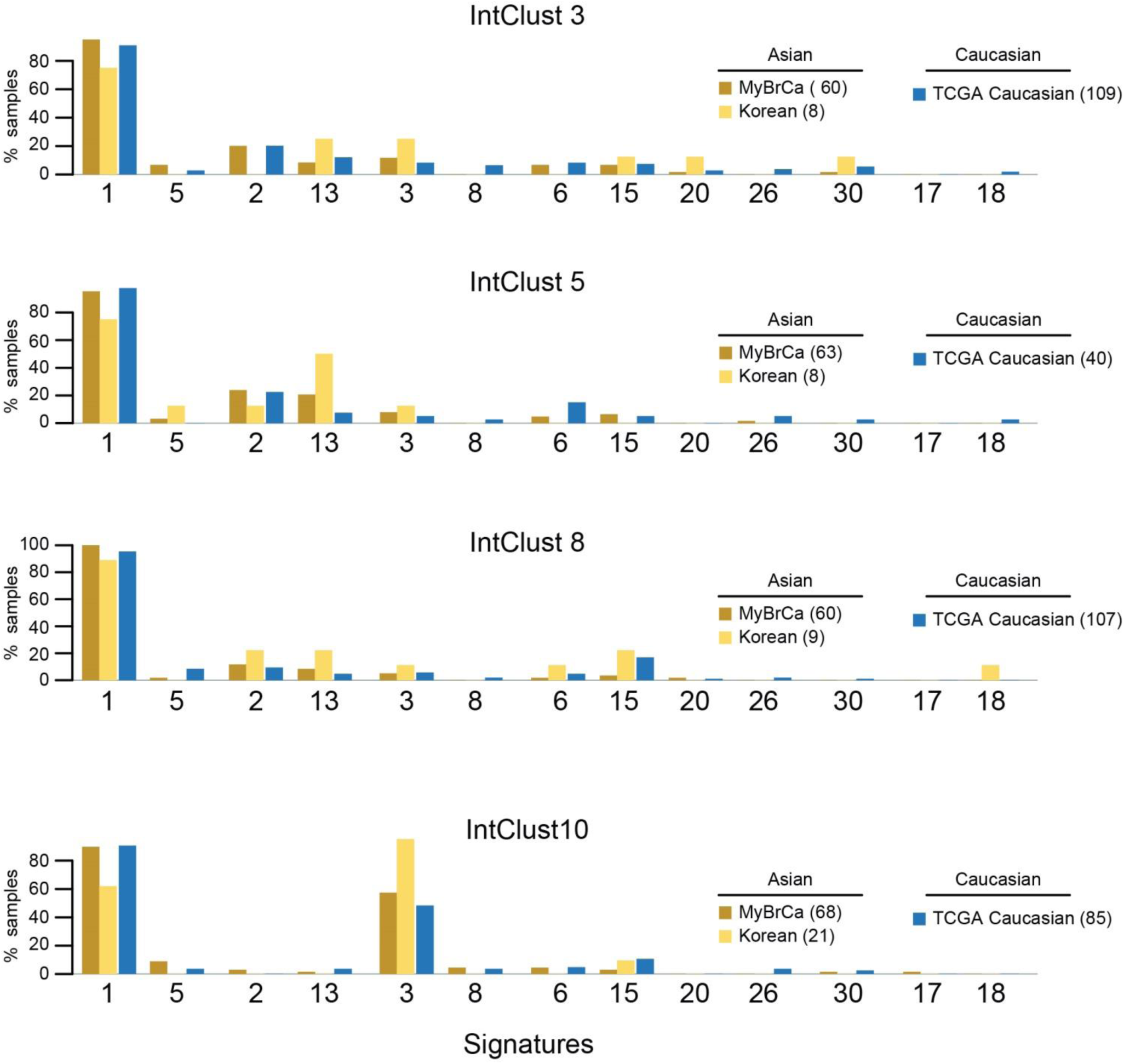
Mutational signatures of Asian and Caucasian breast tumours. Prevalence of mutational signatures in Asian (MyBrCa and Korea) and Caucasian (TCGA Caucasian) tumours, limited to IntClusts 3, 5, 8 and 10.

**SFigure 17:**
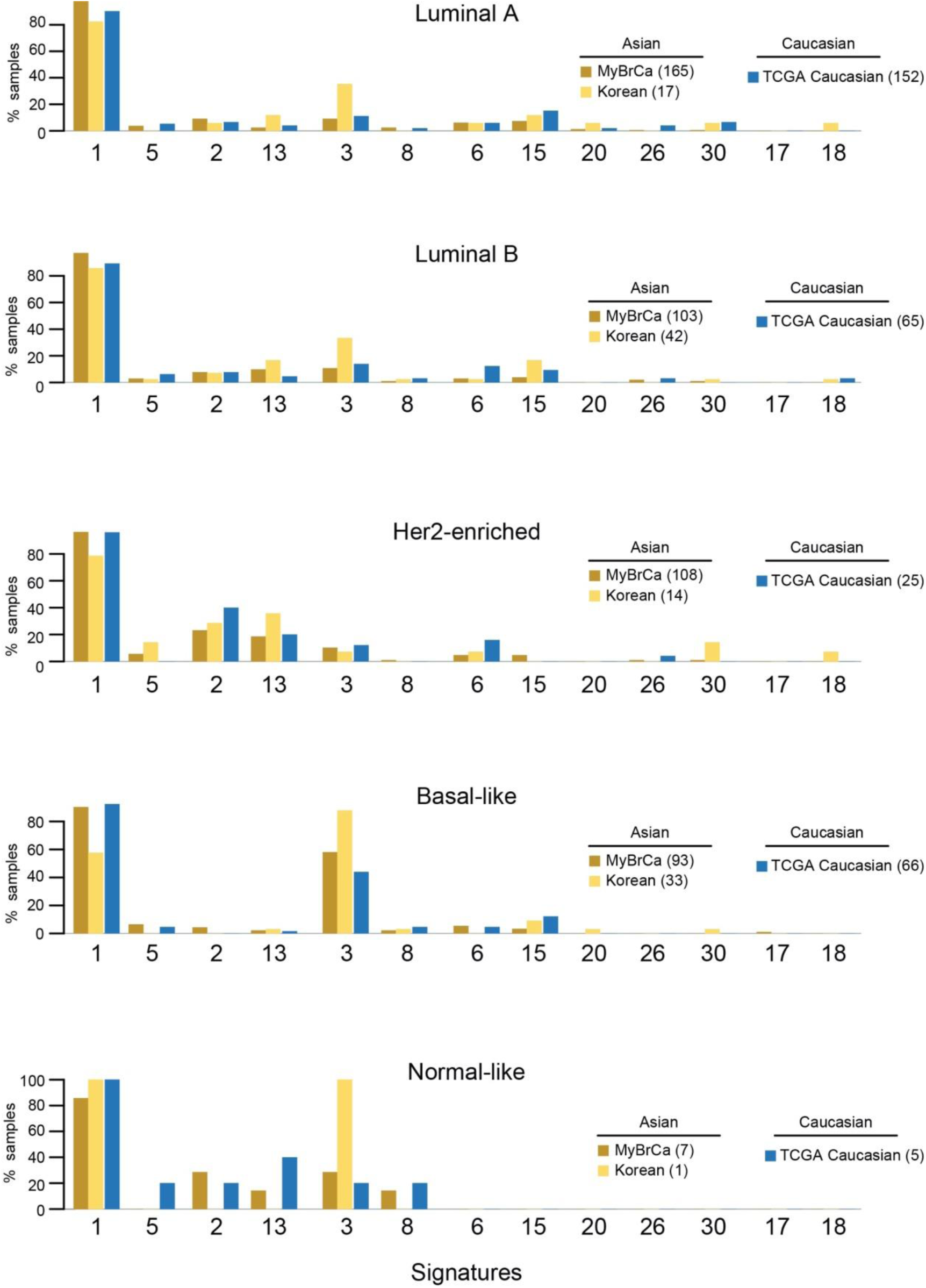
Mutational signatures of Asian and Caucasian breast tumours. Prevalence of mutational signatures in Asian (MyBrCa and Korea) and Caucasian (TCGA Caucasian) breast tumours, limited PAM50 subtypes.

**SFigure 18:**
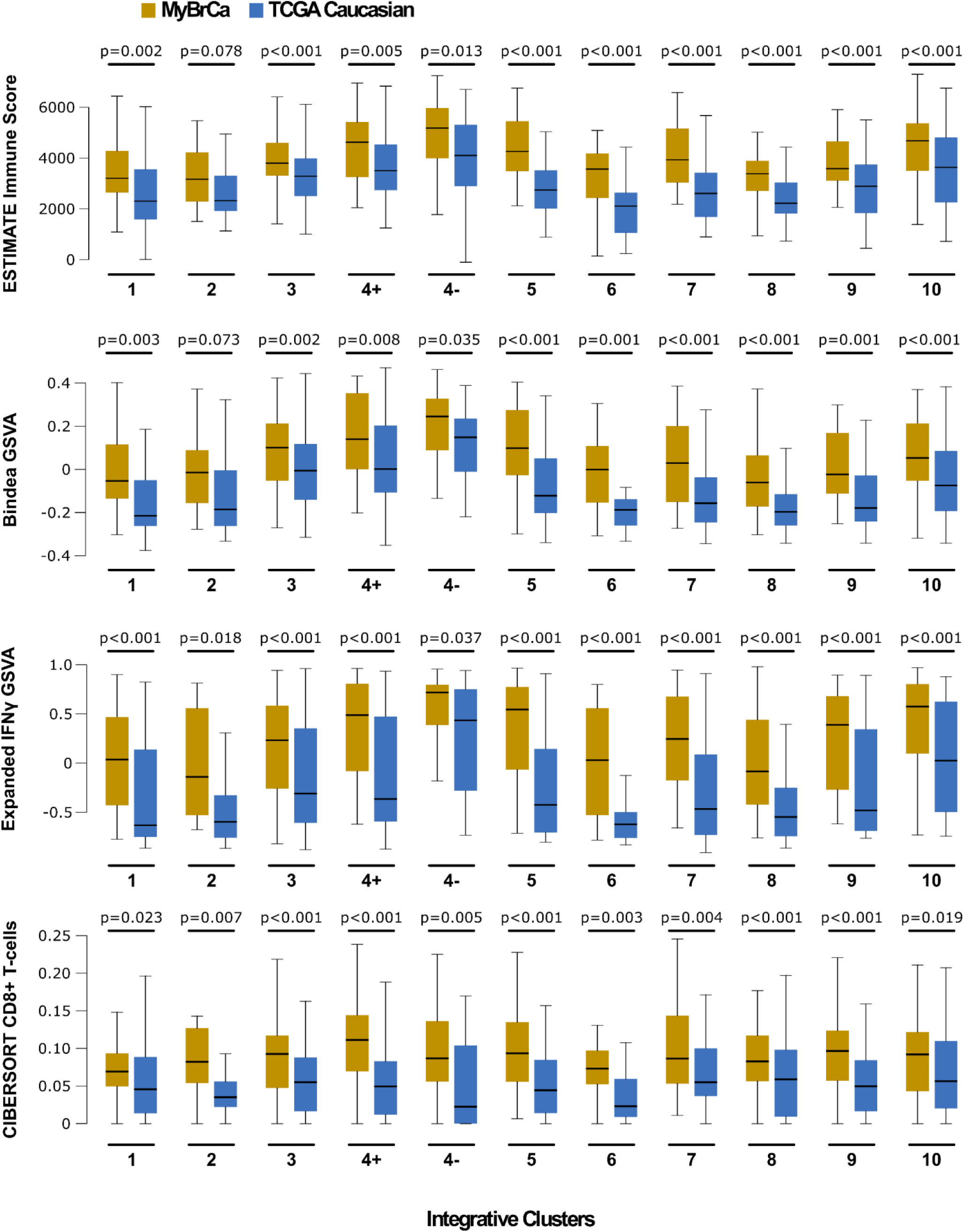
Comparison of immune scores across Integrative Cluster subtypes between MyBrCa samples and Caucasian samples from TCGA. P-values are for *t*-tests between MyBrCa and Caucasian samples for that specific cluster.

**SFigure 19:**
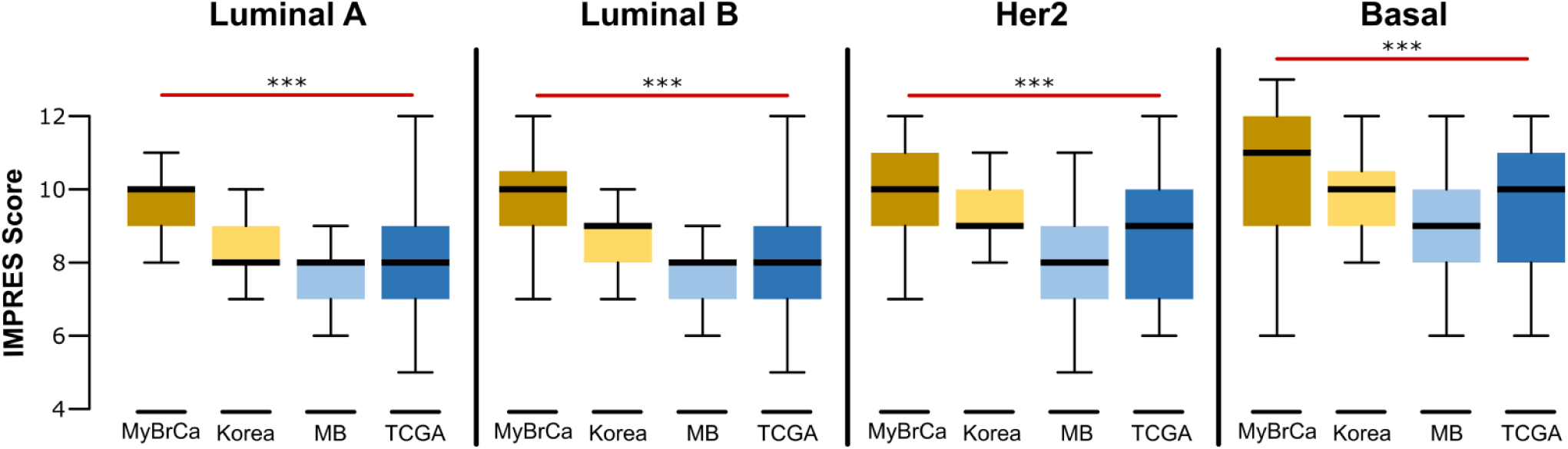
Comparison of IMPRES scores across PAM50 subtypes in the MyBrCa, Korean, METABRIC, and TCGA cohorts. Asterisks indicate significant differences between MyBrCa and TCGA samples from a 2-sided t-test (p<0.001). Outliers not shown.

**SFigure 20:**
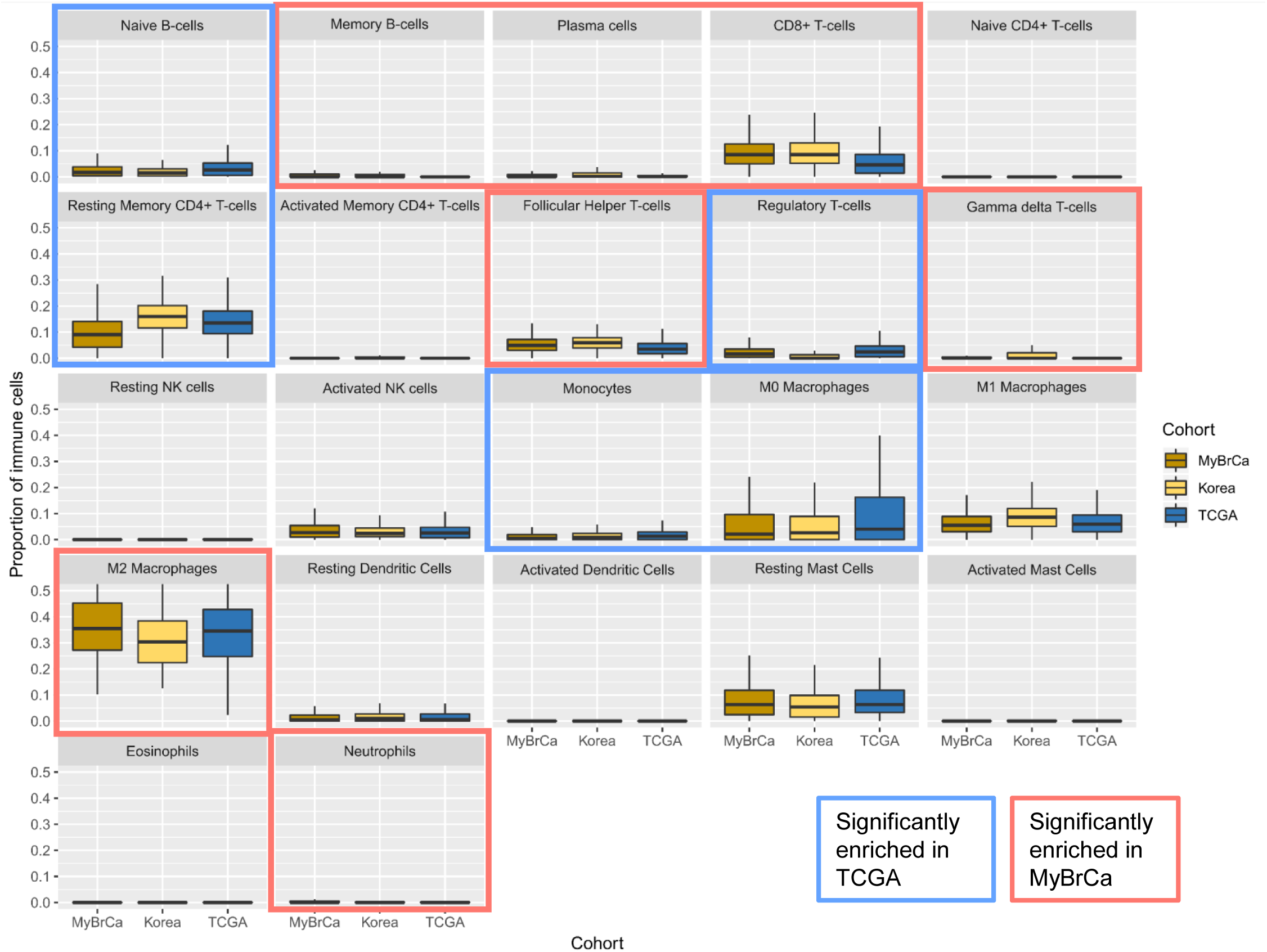
CIBERSORT analysis of MyBrCa, Korean, and TCGA BRCA samples. Highlighted boxes indicate immune cell types that were significantly enriched in either TCGA or MyBrCa relative to the other using a 2-sided t-test (p<0.05). Outliers not shown.

**SFigure 21:**
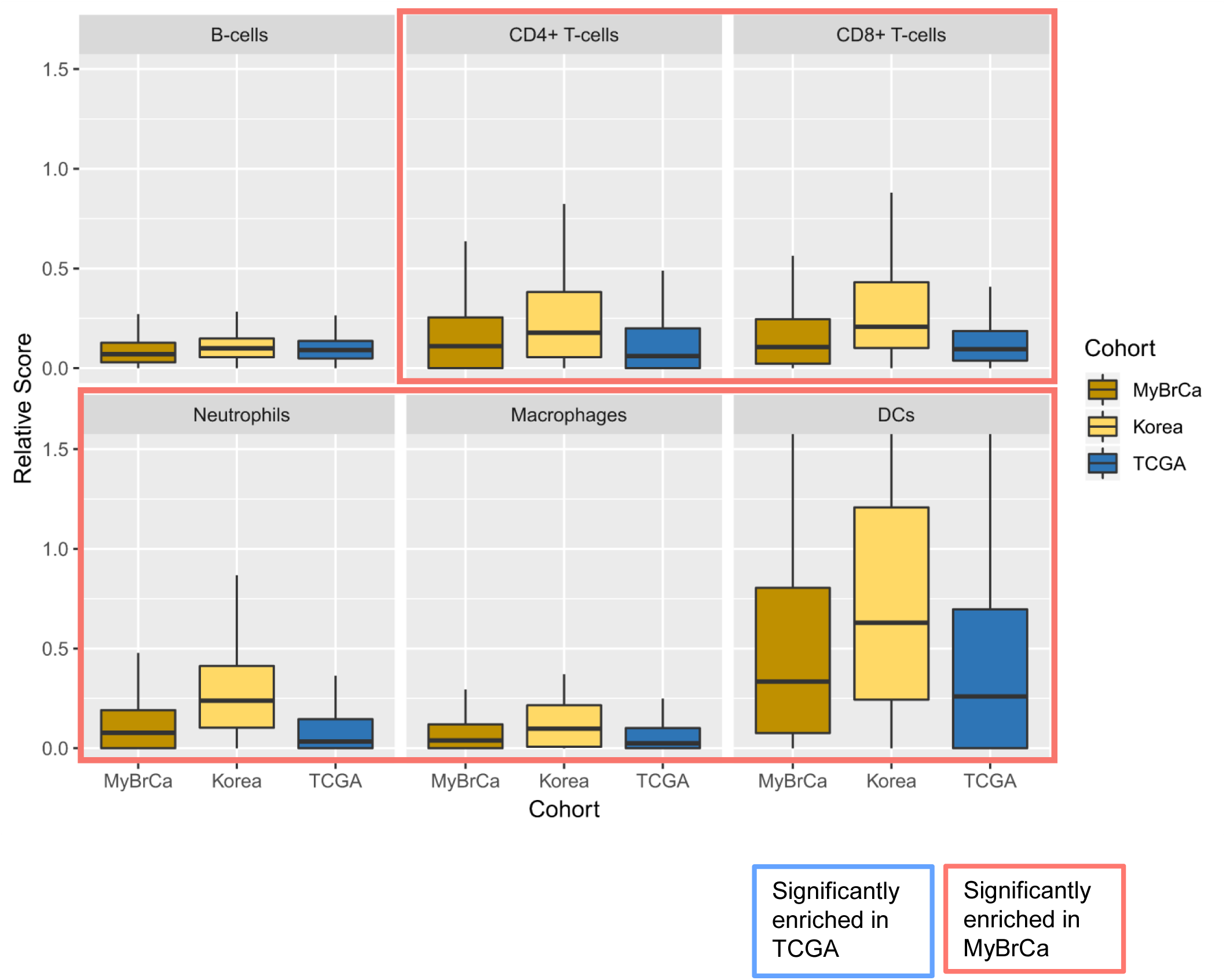
TIMER analysis of MyBrCa, Korean, and TCGA BRCA samples. Highlighted boxes indicate immune cell types that were significantly enriched in either TCGA or MyBrCa relative to the other using a 2-sided *t*-test (p<0.05). Outliers not shown.

**SFigure 22:**
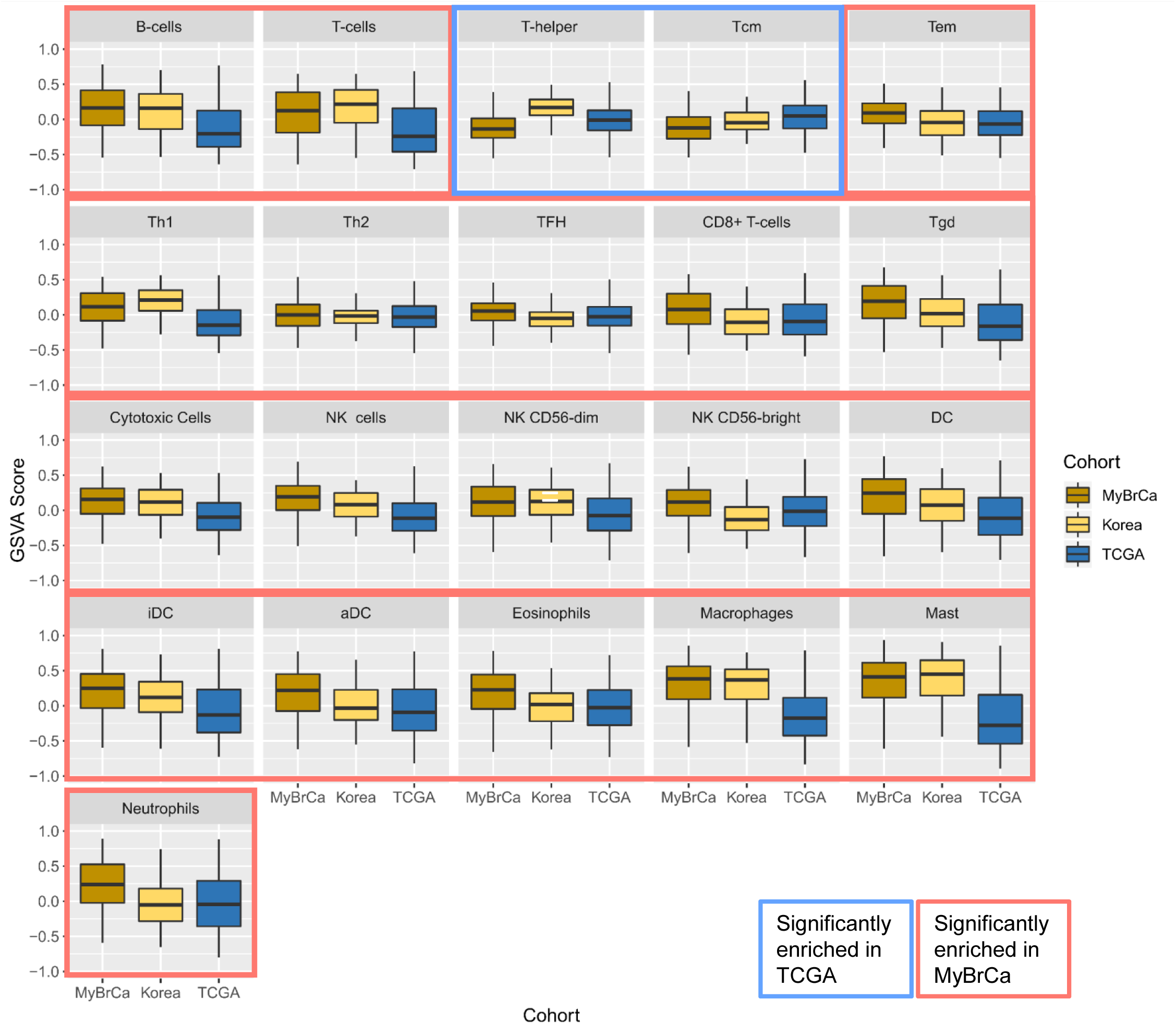
GSVA analysis of MyBrCa, and TCGA BRCA samples using gene sets for individual immune cell types from Bindea et al (2012). Highlighted boxes indicate immune cell types that were significantly enriched in either TCGA or MyBrCa relative to the other using a 2-sided t-test (p<0.05). Outliers not shown.

**SFigure 23:**
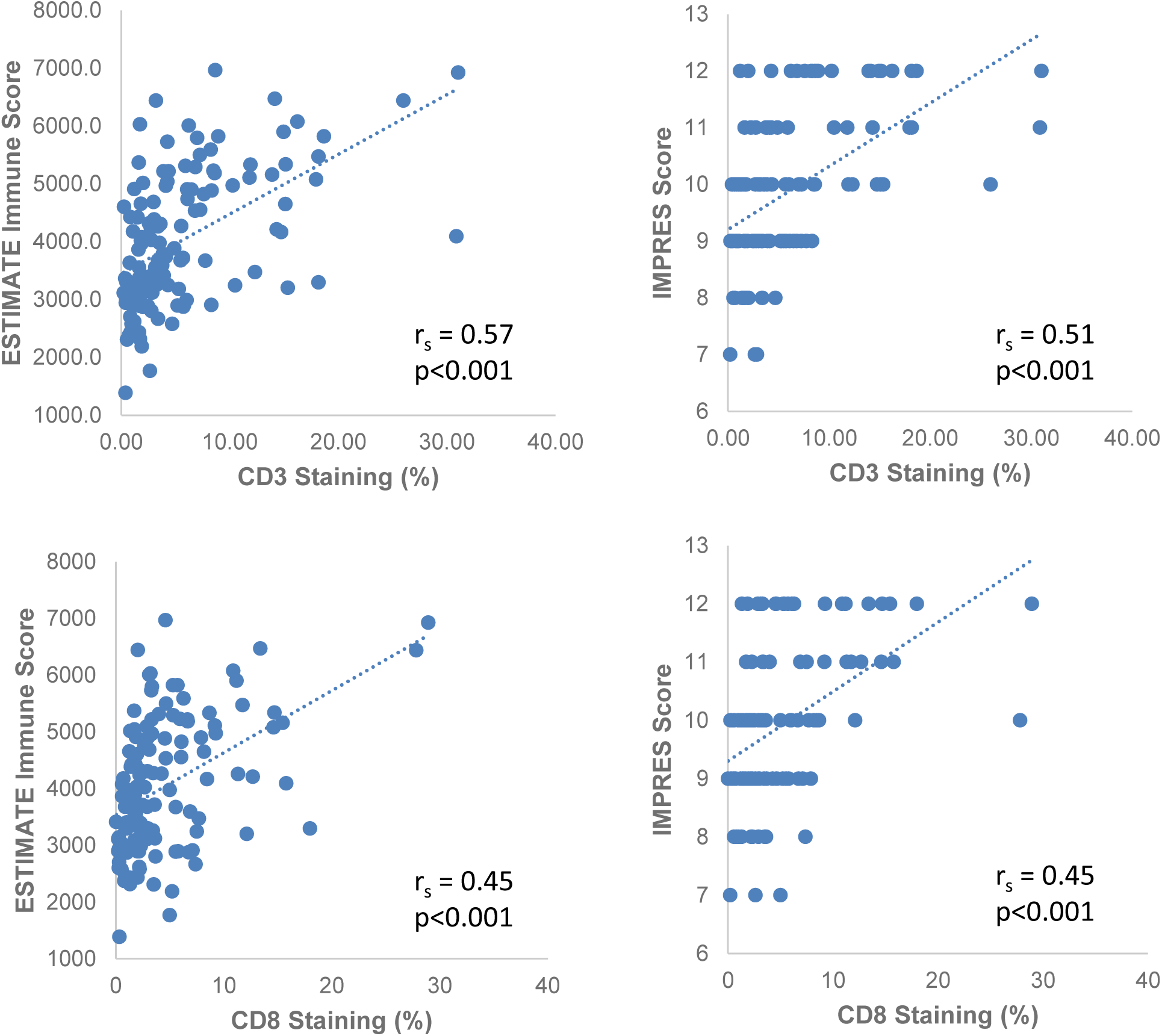
Comparison of ESTIMATE (left) and IMPRES (right) immune scores versus CD3 (top) and CD8 (bottom) IHC staining in the MyBrCa cohort (n=124). Trendlines show linear regression. P-values indicated are for Spearman’s correlation coefficient (r_s_).

**SFigure 24:**
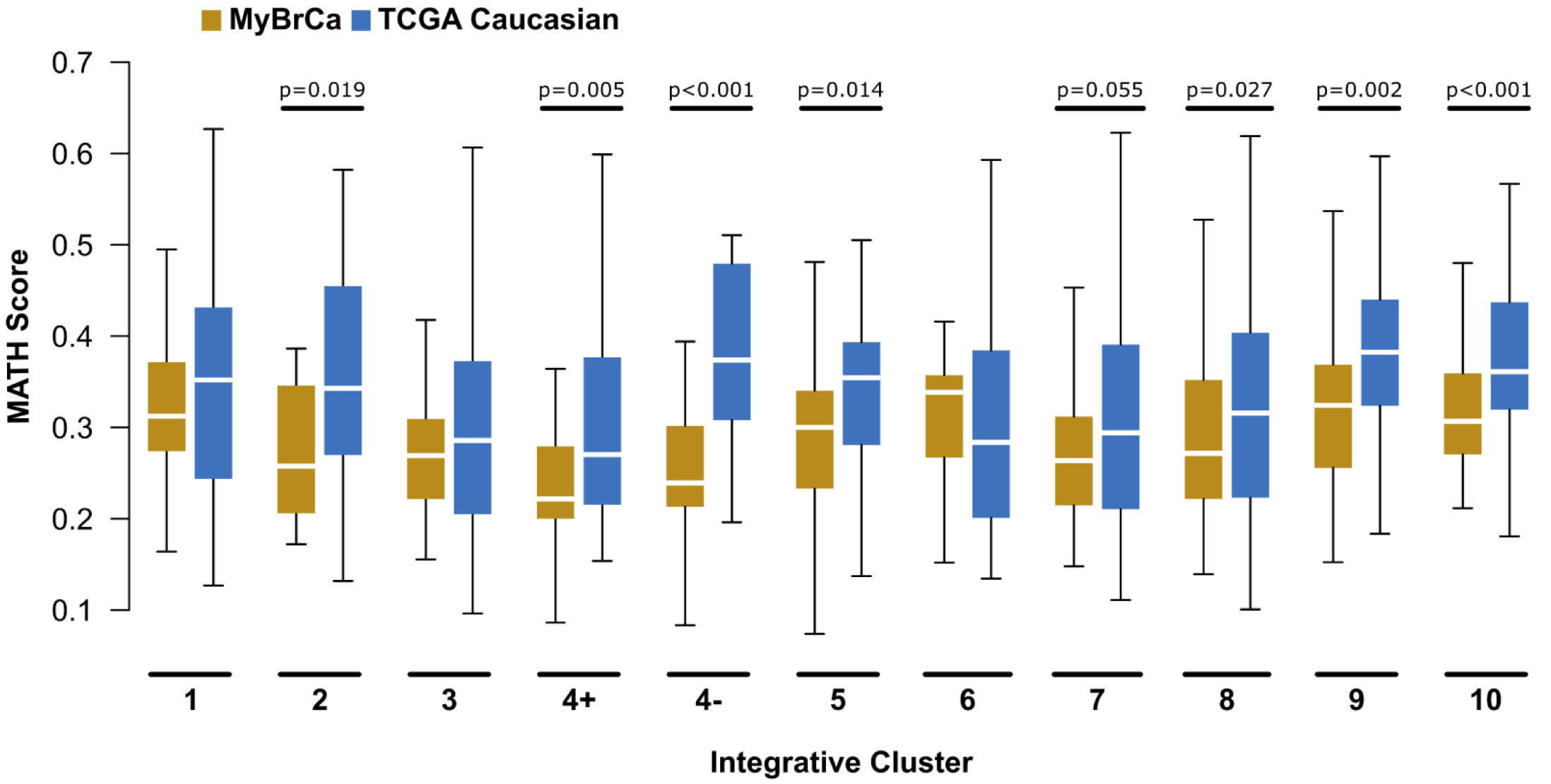
Comparison of tumour heterogeneity between the MyBrCa and TCGA cohorts across Integrative Cluster subtypes. Tumour heterogeneity is quantified here using Mutant Allele Tumour Heterogeneity (MATH) scores.

**SFigure 25.**
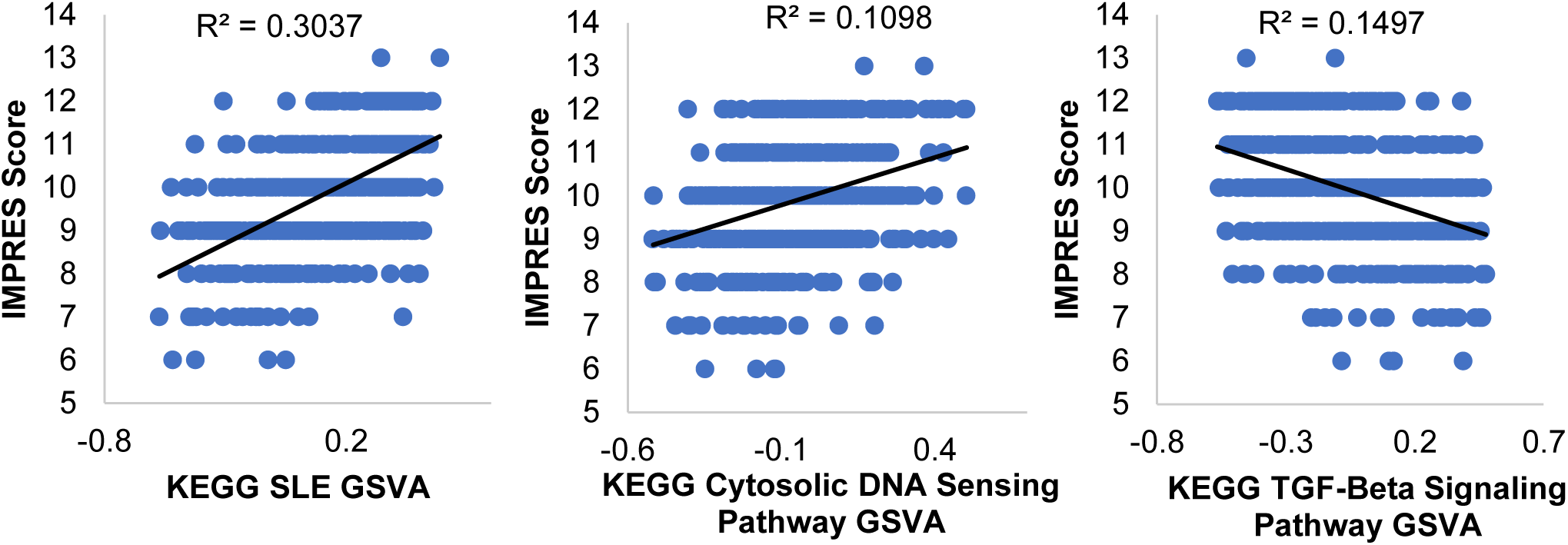
Association of IMPRES scores with GSVA Scores for the KEGG Systemic Lupus Erythematosus pathway, KEGG Cytosolic DNA-sensing pathway, and KEGG TGF-Beta Signaling Pathway. Figures shown are for the MyBrCa cohort samples only.

**SFigure 26:**
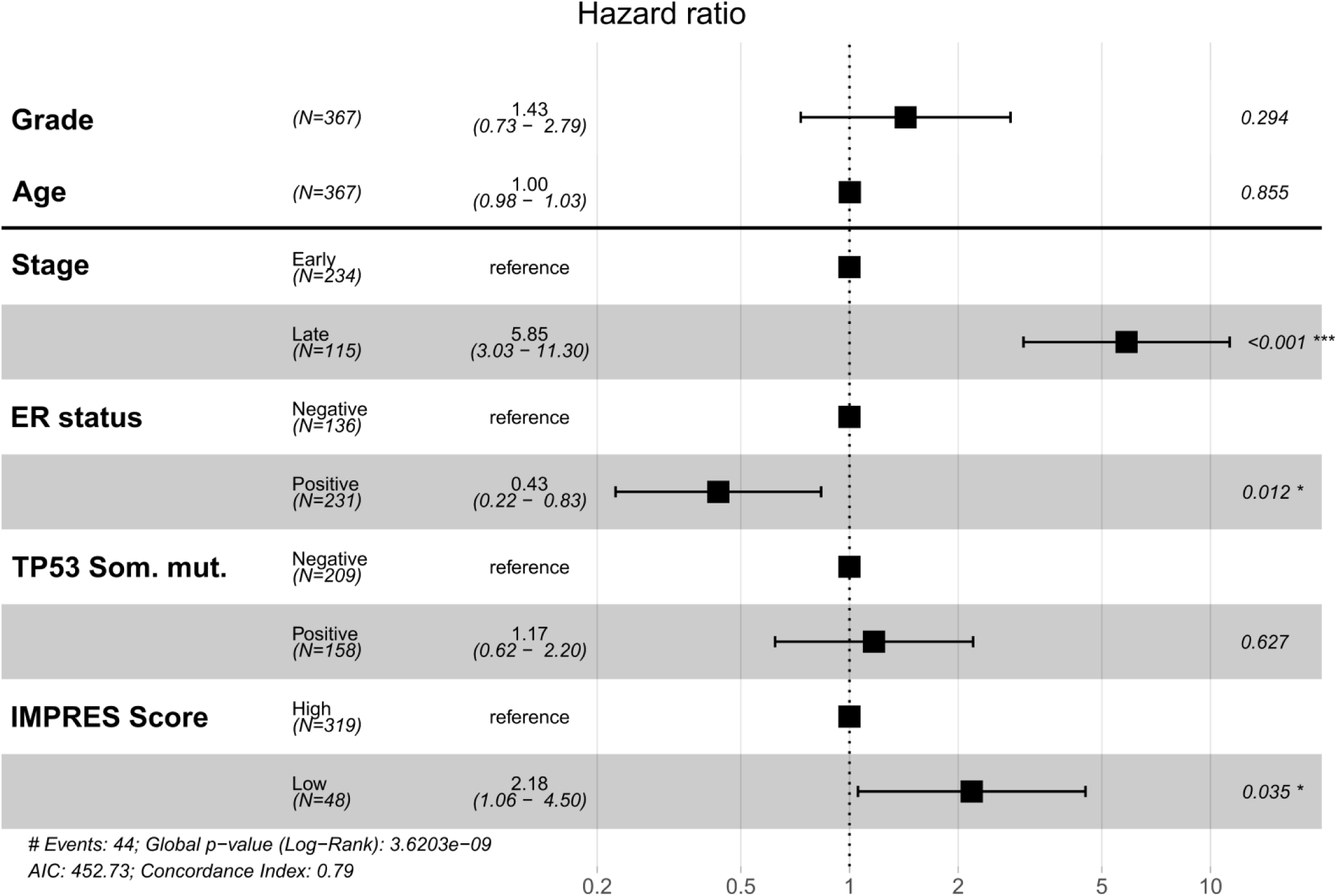
Cox Proportional Hazard model of overall survival for MyBrCa patients. Names of the included variables are bolded on the left, p-values for each individual variable are indicated on the right. Only patients with more than 2 years of follow up data were included (n=367).

**STable 1:**
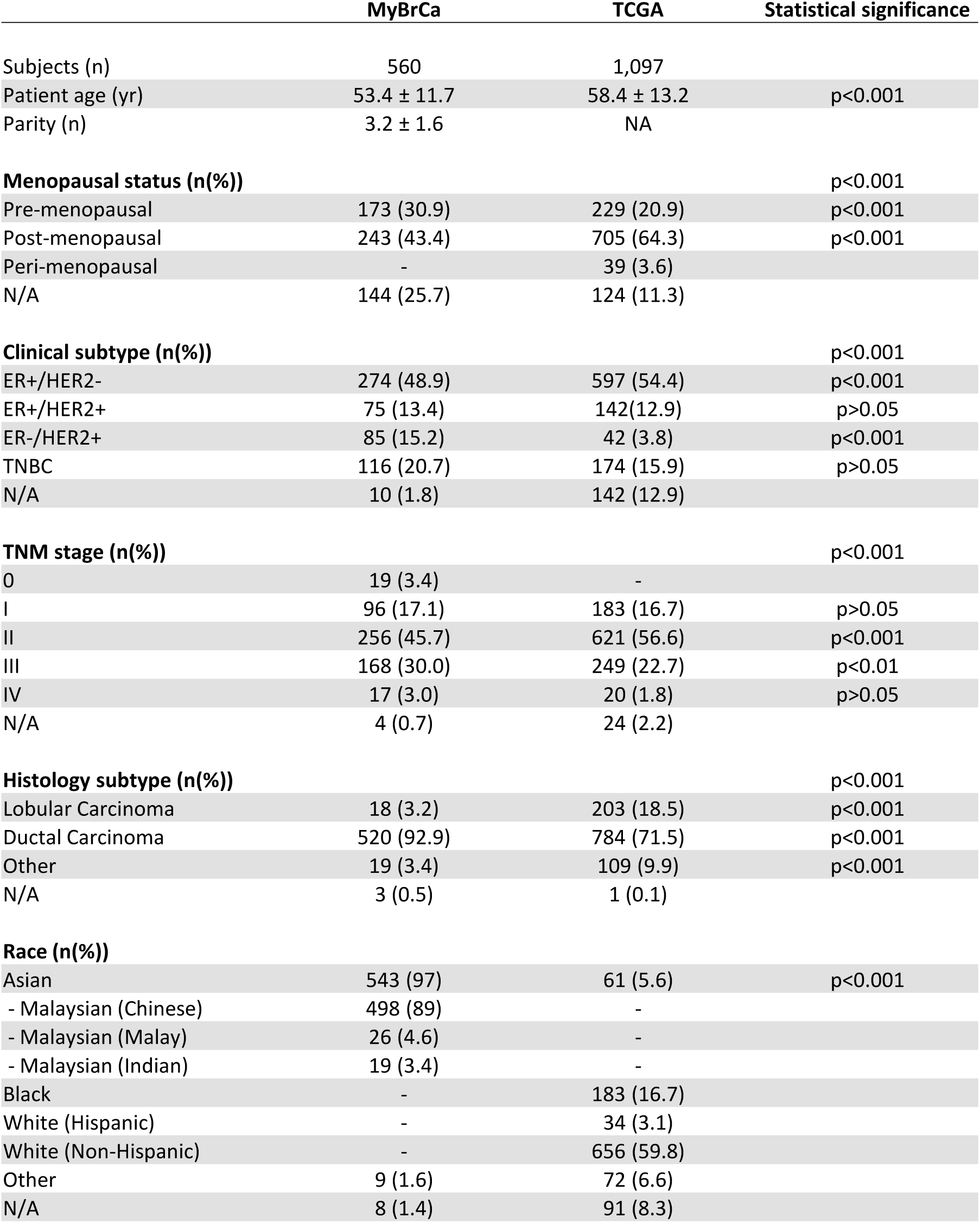
Cohort characteristics. Statistical significance determined with a chi-square test, excluding N/As.

**STable 2:**
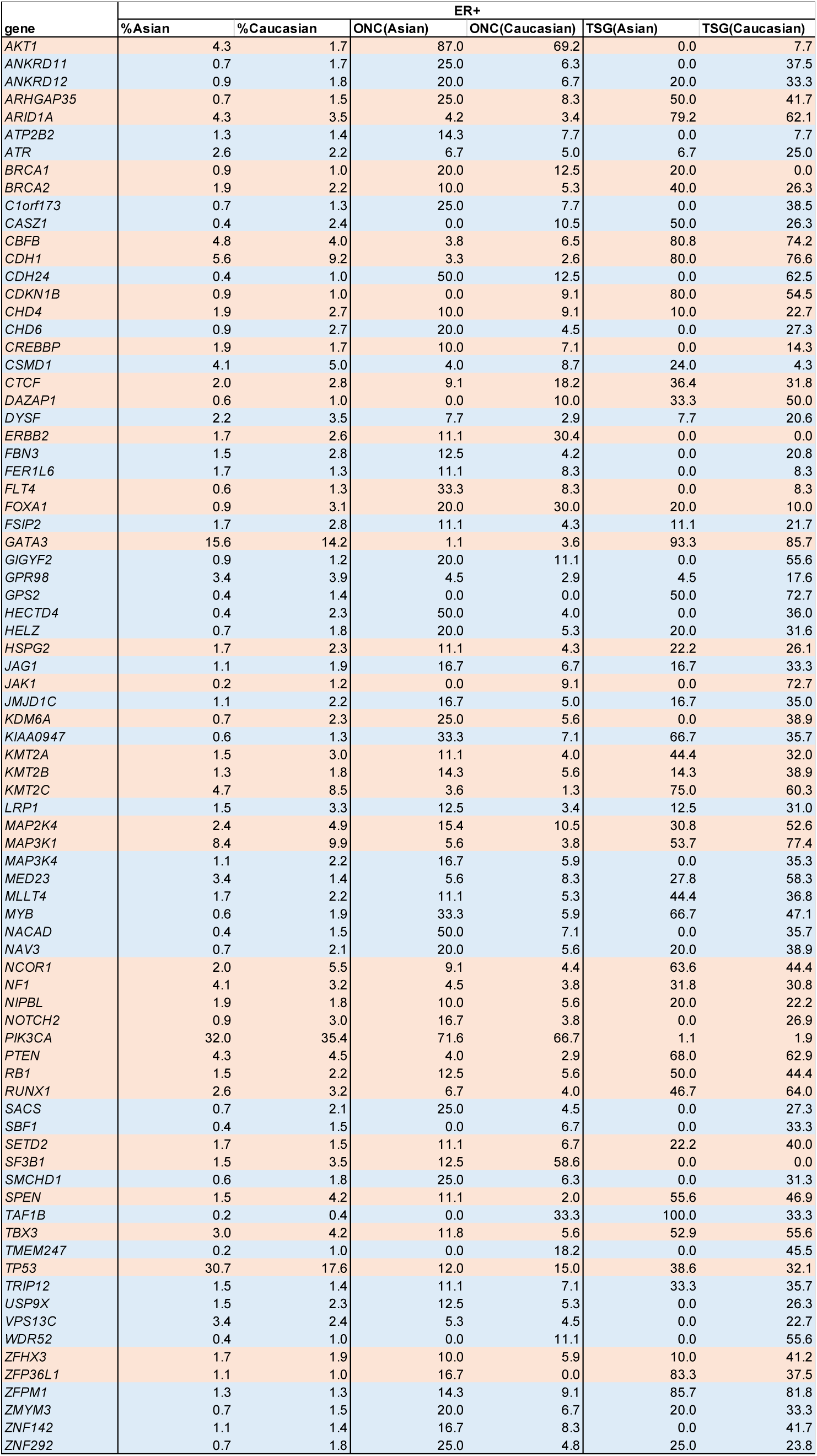
Driver gene analysis on Asian and Caucasian ER+ tumours. Percentage of samples carrying the mutated driver genes, as well as ONC and TSG scores (see Methods) for Asian and Caucasian ER+ tumours. Rows in orange are genes that have been identified as cancer drivers by Integrative Onco Genomics^74^ (see Methods).

**STable 3.**
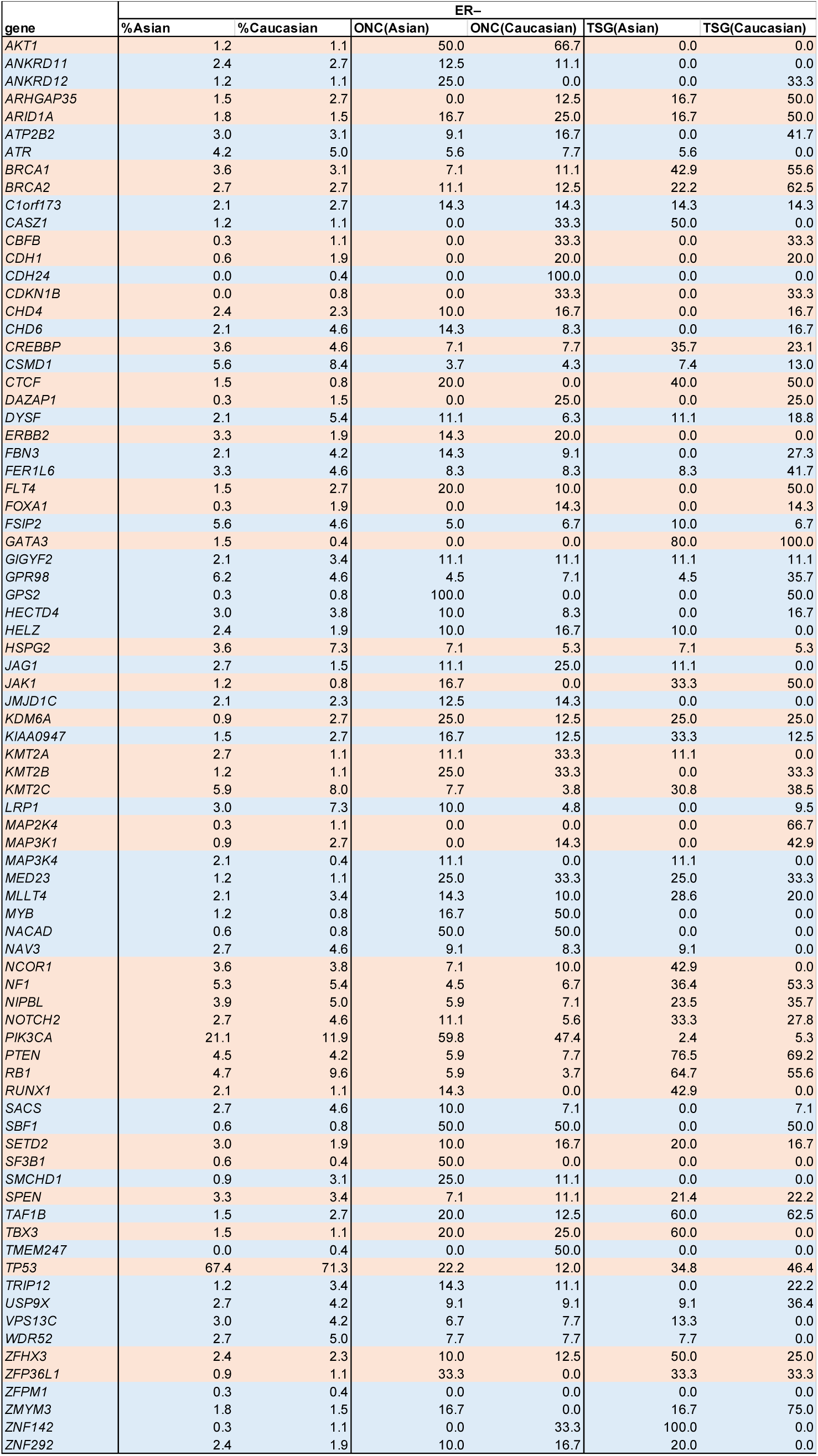
Driver gene analysis on Asian and Caucasian ER– tumours. Percentage of samples carrying the mutated driver genes, as well as ONC and TSG scores (see Methods) for Asian and Caucasian ER– tumours. Rows in orange are genes that have been identified as cancer drivers by Integrative Onco Genomics^74^ (see Methods).

| gene | ER– |  |  |  |  |  |
| --- | --- | --- | --- | --- | --- | --- |
|  | %Asian | %Caucasian | ONC(Asian) | ONC(Caucasian) | TSG(Asian) | TSG(Caucasian) |
| AKT1 | 1.2 | 1.1 | 50.0 | 66.7 | 0.0 | 0.0 |
| ANKRD11 | 2.4 | 2.7 | 12.5 | 11.1 | 0.0 | 0.0 |
| ANKRD12 | 1.2 | 1.1 | 25.0 | 0.0 | 0.0 | 33.3 |
| ARHGAP35 | 1.5 | 2.7 | 0.0 | 12.5 | 16.7 | 50.0 |
| ARID1A | 1.8 | 1.5 | 16.7 | 25.0 | 16.7 | 50.0 |
| ATP2B2 | 3.0 | 3.1 | 9.1 | 16.7 | 0.0 | 41.7 |
| ATR | 4.2 | 5.0 | 5.6 | 7.7 | 5.6 | 0.0 |
| BRCA1 | 3.6 | 3.1 | 7.1 | 11.1 | 42.9 | 55.6 |
| BRCA2 | 2.7 | 2.7 | 11.1 | 12.5 | 22.2 | 62.5 |
| C1orf173 | 2.1 | 2.7 | 14.3 | 14.3 | 14.3 | 14.3 |
| CASZ1 | 1.2 | 1.1 | 0.0 | 33.3 | 50.0 | 0.0 |
| CBFB | 0.3 | 1.1 | 0.0 | 33.3 | 0.0 | 33.3 |
| CDH1 | 0.6 | 1.9 | 0.0 | 20.0 | 0.0 | 20.0 |
| CDH24 | 0.0 | 0.4 | 0.0 | 100.0 | 0.0 | 0.0 |
| CDKN1B | 0.0 | 0.8 | 0.0 | 33.3 | 0.0 | 33.3 |
| CHD4 | 2.4 | 2.3 | 10.0 | 16.7 | 0.0 | 16.7 |
| CHD6 | 2.1 | 4.6 | 14.3 | 8.3 | 0.0 | 16.7 |
| CREBBP | 3.6 | 4.6 | 7.1 | 7.7 | 35.7 | 23.1 |
| CSMD1 | 5.6 | 8.4 | 3.7 | 4.3 | 7.4 | 13.0 |
| CTCF | 1.5 | 0.8 | 20.0 | 0.0 | 40.0 | 50.0 |
| DAZAP1 | 0.3 | 1.5 | 0.0 | 25.0 | 0.0 | 25.0 |
| DYSF | 2.1 | 5.4 | 11.1 | 6.3 | 11.1 | 18.8 |
| ERBB2 | 3.3 | 1.9 | 14.3 | 20.0 | 0.0 | 0.0 |
| FBN3 | 2.1 | 4.2 | 14.3 | 9.1 | 0.0 | 27.3 |
| FER1L6 | 3.3 | 4.6 | 8.3 | 8.3 | 8.3 | 41.7 |
| FLT4 | 1.5 | 2.7 | 20.0 | 10.0 | 0.0 | 50.0 |
| FOXA1 | 0.3 | 1.9 | 0.0 | 14.3 | 0.0 | 14.3 |
| FSIP2 | 5.6 | 4.6 | 5.0 | 6.7 | 10.0 | 6.7 |
| GATA3 | 1.5 | 0.4 | 0.0 | 0.0 | 80.0 | 100.0 |
| GIGYF2 | 2.1 | 3.4 | 11.1 | 11.1 | 11.1 | 11.1 |
| GPR98 | 6.2 | 4.6 | 4.5 | 7.1 | 4.5 | 35.7 |
| GPS2 | 0.3 | 0.8 | 100.0 | 0.0 | 0.0 | 50.0 |
| HECTD4 | 3.0 | 3.8 | 10.0 | 8.3 | 0.0 | 16.7 |
| HELZ | 2.4 | 1.9 | 10.0 | 16.7 | 10.0 | 0.0 |
| HSPG2 | 3.6 | 7.3 | 7.1 | 5.3 | 7.1 | 5.3 |
| JAG1 | 2.7 | 1.5 | 11.1 | 25.0 | 11.1 | 0.0 |
| JAK1 | 1.2 | 0.8 | 16.7 | 0.0 | 33.3 | 50.0 |
| JMJD1C | 2.1 | 2.3 | 12.5 | 14.3 | 0.0 | 0.0 |
| KDM6A | 0.9 | 2.7 | 25.0 | 12.5 | 25.0 | 25.0 |
| KIAA0947 | 1.5 | 2.7 | 16.7 | 12.5 | 33.3 | 12.5 |
| KMT2A | 2.7 | 1.1 | 11.1 | 33.3 | 11.1 | 0.0 |
| KMT2B | 1.2 | 1.1 | 25.0 | 33.3 | 0.0 | 33.3 |
| KMT2C | 5.9 | 8.0 | 7.7 | 3.8 | 30.8 | 38.5 |
| LRP1 | 3.0 | 7.3 | 10.0 | 4.8 | 0.0 | 9.5 |
| MAP2K4 | 0.3 | 1.1 | 0.0 | 0.0 | 0.0 | 66.7 |
| MAP3K1 | 0.9 | 2.7 | 0.0 | 14.3 | 0.0 | 42.9 |
| MAP3K4 | 2.1 | 0.4 | 11.1 | 0.0 | 11.1 | 0.0 |
| MED23 | 1.2 | 1.1 | 25.0 | 33.3 | 25.0 | 33.3 |
| MLLT4 | 2.1 | 3.4 | 14.3 | 10.0 | 28.6 | 20.0 |
| MYB | 1.2 | 0.8 | 16.7 | 50.0 | 0.0 | 0.0 |
| NACAD | 0.6 | 0.8 | 50.0 | 50.0 | 0.0 | 0.0 |
| NAV3 | 2.7 | 4.6 | 9.1 | 8.3 | 9.1 | 0.0 |
| NCOR1 | 3.6 | 3.8 | 7.1 | 10.0 | 42.9 | 0.0 |
| NF1 | 5.3 | 5.4 | 4.5 | 6.7 | 36.4 | 53.3 |
| NIPBL | 3.9 | 5.0 | 5.9 | 7.1 | 23.5 | 35.7 |
| NOTCH2 | 2.7 | 4.6 | 11.1 | 5.6 | 33.3 | 27.8 |
| PIK3CA | 21.1 | 11.9 | 59.8 | 47.4 | 2.4 | 5.3 |
| PTEN | 4.5 | 4.2 | 5.9 | 7.7 | 76.5 | 69.2 |
| RB1 | 4.7 | 9.6 | 5.9 | 3.7 | 64.7 | 55.6 |
| RUNX1 | 2.1 | 1.1 | 14.3 | 0.0 | 42.9 | 0.0 |
| SACS | 2.7 | 4.6 | 10.0 | 7.1 | 0.0 | 7.1 |
| SBF1 | 0.6 | 0.8 | 50.0 | 50.0 | 0.0 | 50.0 |
| SETD2 | 3.0 | 1.9 | 10.0 | 16.7 | 20.0 | 16.7 |
| SF3B1 | 0.6 | 0.4 | 50.0 | 0.0 | 0.0 | 0.0 |
| SMCHD1 | 0.9 | 3.1 | 25.0 | 11.1 | 0.0 | 0.0 |
| SPEN | 3.3 | 3.4 | 7.1 | 11.1 | 21.4 | 22.2 |
| TAF1B | 1.5 | 2.7 | 20.0 | 12.5 | 60.0 | 62.5 |
| TBX3 | 1.5 | 1.1 | 20.0 | 25.0 | 60.0 | 0.0 |
| TMEM247 | 0.0 | 0.4 | 0.0 | 50.0 | 0.0 | 0.0 |
| TP53 | 67.4 | 71.3 | 22.2 | 12.0 | 34.8 | 46.4 |
| TRIP12 | 1.2 | 3.4 | 14.3 | 11.1 | 0.0 | 22.2 |
| USP9X | 2.7 | 4.2 | 9.1 | 9.1 | 9.1 | 36.4 |
| VPS13C | 3.0 | 4.2 | 6.7 | 7.7 | 13.3 | 0.0 |
| WDR52 | 2.7 | 5.0 | 7.7 | 7.7 | 7.7 | 0.0 |
| ZFHX3 | 2.4 | 2.3 | 10.0 | 12.5 | 50.0 | 25.0 |
| ZFP36L1 | 0.9 | 1.1 | 33.3 | 0.0 | 33.3 | 33.3 |
| ZFPM1 | 0.3 | 0.4 | 0.0 | 0.0 | 0.0 | 0.0 |
| ZMYM3 | 1.8 | 1.5 | 16.7 | 0.0 | 16.7 | 75.0 |
| ZNF142 | 0.3 | 1.1 | 0.0 | 33.3 | 100.0 | 0.0 |
| ZNF292 | 2.4 | 1.9 | 10.0 | 16.7 | 20.0 | 0.0 |

**STable 4:** Pathway analysis of MyBrCa versus TCGA tumours. Top 15 most differentially enriched pathways between the MyBrCa and TCGA cohorts as indicated by Gene Set Enrichment Analysis of MSigDB hallmark pathways, ranked by absolute normalized enrichment score (|NES|). Immune system-related gene sets are highlighted in blue.

| Rank | Hallmark Pathway | \|NES\| | p-val | FDR | Higher in |
| --- | --- | --- | --- | --- | --- |
| 1 | Mitotic Spindle | 2.061 | 0 | 0.012 | TCGA |
| 2 | Estrogen Response Early | 2.016 | 0 | 0.009 | TCGA |
| 3 | UV Response Down | 1.953 | 0 | 0.018 | TCGA |
| 4 | Allograft Rejection | 1.928 | 0.004 | 0.088 | MyBrCa |
| 5 | TGF Beta Signaling | 1.818 | 0.011 | 0.041 | TCGA |
| 6 | Notch Signaling | 1.762 | 0.007 | 0.051 | TCGA |
| 7 | Apical Surface | 1.753 | 0.004 | 0.047 | TCGA |
| 8 | Wnt Beta Catenin Signaling | 1.716 | 0.012 | 0.051 | TCGA |
| 9 | Complement | 1.709 | 0.019 | 0.289 | MyBrCa |
| 10 | Oxidative Phosphorylation | 1.578 | 0.092 | 0.418 | MyBrCa |
| 11 | Inflammatory Response | 1.565 | 0.100 | 0.339 | MyBrCa |
| 12 | Interferon Gamma Response | 1.551 | 0.113 | 0.292 | MyBrCa |
| 13 | Interferon Alpha Response | 1.542 | 0.117 | 0.253 | MyBrCa |
| 14 | Hedgehog Signaling | 1.526 | 0.047 | 0.139 | TCGA |
| 15 | Adipogenesis | 1.525 | 0.061 | 0.239 | MyBrCa |

**Stable 5:** Regression analysis of IMPRES score in the MyBrCa cohort (n=340) using a multivariable linear model across available clinical and demographic data. Asterisks indicate the level of significance (Pr(>|t|)) for each variable (*< 0.05; **<0.01; ***<0.001).

|  | Variable | Est. Coefficient | Std. Error | t-value | Pr(>\|t\|) | Sig. |
| --- | --- | --- | --- | --- | --- | --- |
| **Integrative Cluster Subtype**<br>(Relative to IntClust 3) | IntClust 1 | -0.131 | 0.341 | -0.383 | 0.702 |  |
|  | IntClust 2 | 0.281 | 0.464 | 0.606 | 0.545 |  |
|  | IntClust 4+ | 0.329 | 0.319 | 1.032 | 0.303 |  |
|  | **IntClust 4-** | **1.013** | **0.348** | **2.916** | **0.004** | **\*\*** |
|  | IntClust 5 | 0.335 | 0.304 | 1.101 | 0.272 |  |
|  | IntClust 6 | 0.091 | 0.421 | 0.215 | 0.830 |  |
|  | **IntClust 7** | **0.658** | **0.310** | **2.126** | **0.034** | **\*** |
|  | IntClust 8 | 0.012 | 0.291 | 0.04 | 0.968 |  |
|  | **IntClust 9** | **0.644** | **0.318** | **2.029** | **0.043** | **\*** |
|  | **IntClust 10** | **1.007** | **0.312** | **3.227** | **0.001** | **\*\*** |
| **Histological subtype**<br>(Relative to DCIS) | Histology - IDC | -0.198 | 0.587 | -0.338 | 0.735 |  |
|  | Histology - ILC | -0.701 | 0.768 | -0.913 | 0.362 |  |
|  | Histology - Other | -0.560 | 0.978 | -0.572 | 0.567 |  |
| **Clinical metrics** | Grade | 0.185 | 0.149 | 1.242 | 0.215 |  |
|  | Stage (AJCC6) | 0.108 | 0.105 | 1.028 | 0.305 |  |
|  | Size | -0.019 | 0.051 | -0.383 | 0.702 |  |
|  | Tumor Content | 0.000 | 0.004 | -0.005 | 0.996 |  |
|  | Neoadjuvant treatment | -0.309 | 0.413 | -0.748 | 0.455 |  |
| **Physical metrics** | **Age** | **-0.015** | **0.008** | **-2.012** | **0.045** | **\*** |
|  | Weight | -0.052 | 0.098 | -0.533 | 0.594 |  |
|  | Height | 3.350 | 7.421 | 0.451 | 0.652 |  |
|  | BMI | 0.118 | 0.243 | 0.485 | 0.628 |  |
| **Demographic factors**<br><sup>1</sup>Relative to Chinese<br><sup>2</sup>Relative to Income <5,000 | Ethnicity – Indian<sup>1</sup> | -0.214 | 0.441 | -0.486 | 0.628 |  |
|  | Ethnicity - Malay<sup>1</sup> | -0.238 | 0.382 | -0.621 | 0.535 |  |
|  | Ethnicity - Other<sup>1</sup> | -0.250 | 0.530 | -0.472 | 0.637 |  |
|  | Age of menarche | 0.057 | 0.055 | 1.041 | 0.299 |  |
|  | Family history of cancer | 0.173 | 0.181 | 0.953 | 0.341 |  |
|  | Income >10,000<sup>2</sup> | 0.108 | 0.230 | 0.471 | 0.638 |  |
|  | Income 5-10,000<sup>2</sup> | 0.042 | 0.180 | 0.231 | 0.817 |  |
|  | Education | -0.167 | 0.130 | -1.289 | 0.198 |  |
|  | Regular alcohol drinker | 0.064 | 0.202 | 0.316 | 0.752 |  |

Adjusted R-squared: 0.0674
Model p-value: 0.0047

**STable 6:**
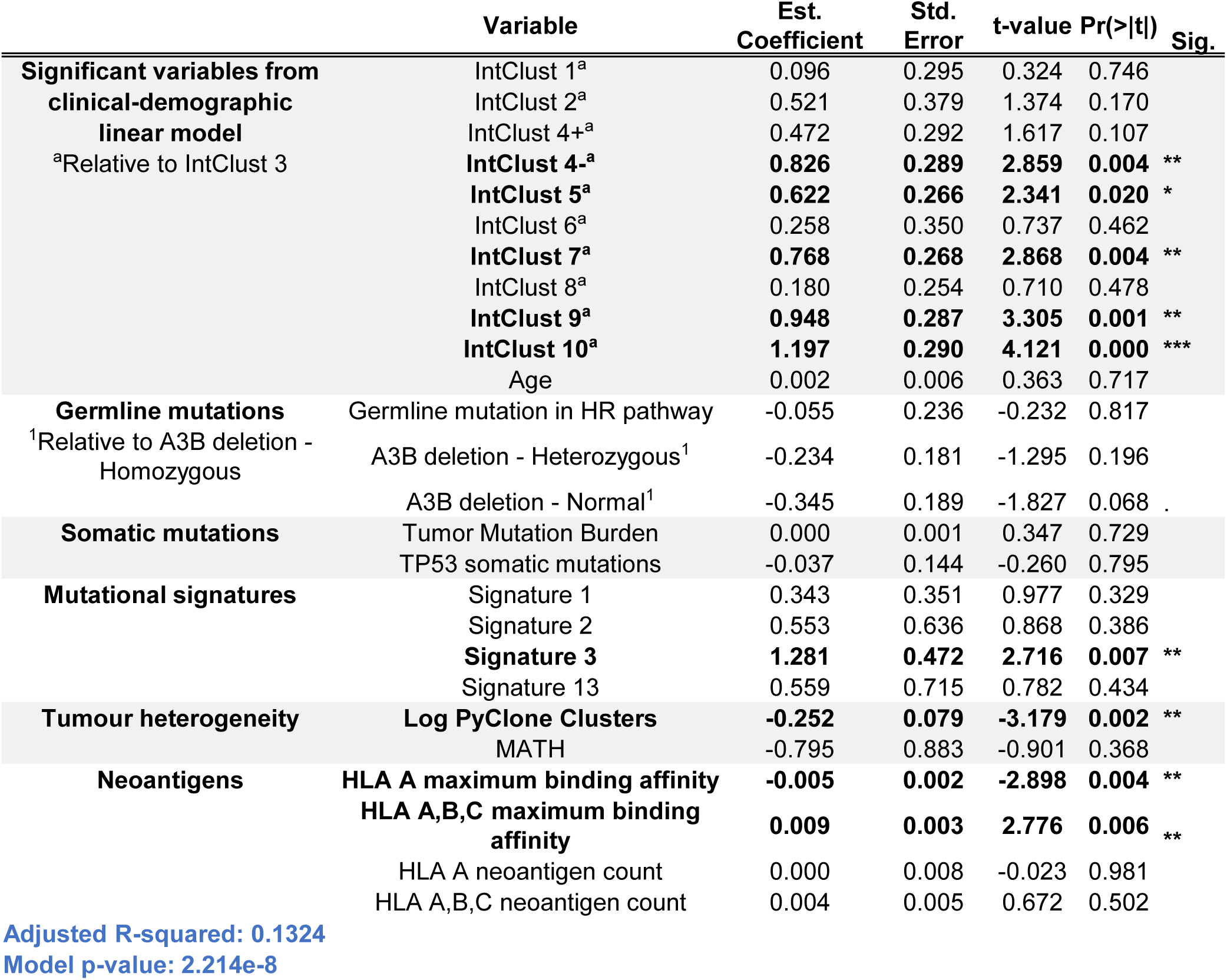
Linear regression analysis of IMPRES score in the MyBrCa cohort (n=340) using available molecular data, adjusting for age and subtype. Asterisks indicate the level of significance (Pr(>|t|)) for each variable (*< 0.05; **<0.01; ***<0.001).

**STable 7:** Pathway analysis of IMPRES high- versus low-scoring tumours. Pathway analysis comparing MyBrCa samples in the top quartile of IMPRES scores to samples in the bottom quartile reveals significant differences in the SLE pathway, cytosolic DNA-sensing pathway and TFG-Beta signaling pathway between the two groups.

| KEGG Pathway | \|NES\| | FDR | Enriched in |
| --- | --- | --- | --- |
| Systemic Lupus Erythematosus | 2.29 | 0 | Top Quartile |
| Cytosolic DNA-sensing Pathway | 2.02 | $3 \times 10^{-4}$ | Top Quartile |
| TGF-Beta Signaling Pathway | 1.79 | 0.04 | Bottom Quartile |

